# Targeting AXL Overcomes Adaptive Resistance to KRAS Inhibition in KRAS-Driven Cancer

**DOI:** 10.64898/2026.08.02.742234

**Authors:** Fredrik I. Thege, Annaliese Kramer, Norbert Kreisz, Intisar Salim, Etsehiwot Girum Girma, Linara Welte, Elizabeth Adams, Ashton Pluchinsky, Elijah Kirschstein, Olivia Harder, Amber Hoskins, Ashwath Seetharaman, Kimal Rajapakshe, Yuki Makino, Andrew J. Gunderson, Sonja M. Wörmann, Anirban Maitra

## Abstract

Direct KRAS inhibitors have established mutant KRAS as a clinically actionable target, yet adaptive resistance remains a major barrier to durable responses. To identify therapeutically actionable resistance mechanisms, we performed an unbiased *in vivo* CRISPR activation screen in an autochthonous lung adenocarcinoma model, identifying the receptor tyrosine kinase AXL as a dominant adaptive resistance driver. Pharmacologic AXL inhibition enhanced the efficacy of both allele-specific inhibition and the RAS(ON) multi-selective inhibitor daraxonrasib across lung and pancreatic cancer models, resulting in deeper and more durable suppression of MAPK signaling and improved tumor control. Beyond tumor-intrinsic effects, combined KRAS and AXL inhibition remodeled the tumor immune microenvironment, promoting an IFNγ-responsive program, increased recruitment of cytotoxic T cells and sensitization to FAS-mediated apoptosis. Collectively, our findings identify AXL as a convergence point for adaptive resistance to KRAS inhibition and provide a mechanistically informed combination strategy to extend the durability of KRAS-directed therapies.

**Statement of Significance:** An unbiased in vivo functional (CRISPR activation) screen identifies AXL as a convergence point for adaptive resistance to KRAS inhibition. By integrating adaptive response to KRAS inhibition with anti-tumor immunity, AXL represents a mechanistically actionable vulnerability whose inhibition deepens and prolongs responses to both allele-specific and pan-KRAS-targeted therapies.

## Introduction

Direct pharmacologic inhibition of mutant KRAS has transformed one of the longest-standing challenges in cancer therapeutics into a clinical reality. The clinical success of the KRAS^G12C^ inhibitors sotorasib and adagrasib, together with the rapid development of next-generation allele-specific inhibitors and RAS(ON) multi-selective inhibitors including daraxonrasib (RMC-6236), has established mutant KRAS as a tractable therapeutic target across multiple cancer types(1–5). Yet, despite unprecedented progress, durable responses remain uncommon, highlighting adaptive resistance as the principal obstacle to maximizing the therapeutic potential of KRAS inhibition.

Mechanisms of resistance to KRAS inhibitors are increasingly recognized to arise through both genetic evolution and non-genetic adaptive remodeling. Acquired alterations involving KRAS, NRAS, BRAF, and other MAPK pathway components restore ERK signaling during prolonged treatment(6,7). However, multiple studies have demonstrated that rapid transcriptional and signaling adaptation frequently precedes these genomic events, allowing tumor cells to survive initial therapeutic pressure and providing a substrate for subsequent genetic evolution(8,9). These observations suggest that the earliest determinants of therapeutic failure are likely governed by dynamic signaling networks rather than irreversible genomic alterations.

Among these adaptive programs, reactivation of receptor tyrosine kinase (RTK) signaling has emerged as one of the most consistent features of resistance across distinct classes of KRAS inhibitors(3,6,10). Multiple RTKs (including EGFR, FGFR, ERBB2, and MET) have been implicated in adaptive therapy escape. Despite these advances, it remains unclear whether shared signaling nodes integrate these diverse adaptive responses and how adaptive signaling interfaces with the tumor immune microenvironment, which is increasingly recognized as a critical determinant of response to KRAS-targeted therapies.

To address this question, we performed an unbiased in vivo CRISPR activation screen in an autochthonous model of KRAS-mutant lung adenocarcinoma, enabling systematic identification of resistance mechanisms operating within the native tumor ecosystem. We identify the receptor tyrosine kinase AXL as a dominant mediator of adaptive resistance across distinct classes of KRAS inhibitors. Our findings establish a mechanistic framework for rational KRAS-directed combination therapies and support concurrent KRAS and AXL inhibition as a strategy to improve the durability of KRAS-targeted treatment.

## Results

### Autochthonous in vivo CRISPR activation drug screening identifies AXL as a candidate for combinatorial KRAS inhibition

To identify candidates for combinatorial targeting with KRAS inhibition, we performed autochthonous in vivo CRISPRa lung tumor screening using our recently published FiCASCan (*Focused in vivo CRISPR Activation Screening for Cancer*) platform(11,12). Our genetically engineered LSL-SAM mouse model allows for programmable gene activation in vivo and, when combined with conditional oncogenic Kras^LSL-G12D^ and P53 loss (P53^F/F^), constitutes the CRISPRa-competent “PPKS” (Trp53F/F; Kras^LSL-G12D^; Rosa26^LSL-SAM^) mouse tumor model(11). FiCASCan works by introducing pooled Cre/sgRNA-encoding lentivirus through nasal instillation in PPKS mice to initiate lung tumor formation and concurrent gene activation in vivo. Here, we use FiCASCan in conjunction with KRAS inhibitor treatment (KRASi; MRTX1133) to select guides that enable tumor lesions to persist and/or expand despite treatment. MRTX1133 is a non-covalent KRAS(OFF) inhibitor (PMID:34889605) that has shown efficacy in preclinical models(13–15). To this end, we generated lung tumors by treating thirteen PPKS mice with 1e8 TU of pooled lentivirus encoding for the Onco1 sgRNA(MS2) library (Figure 1A)(12), which targets 442 of the most commonly altered genes across human adenocarcinoma with 4 guides per transcription start site, in addition to 100 non-targeted guides (1976 guides in total). Six weeks after tumor induction, we initiated treatment with KRASi or vehicle, once daily, for four weeks (Figure 1B). The lungs were then harvested, gDNA isolated, and the sgRNA distribution was determined by NGS (Figure S1A-D). We next identified genes enriched in the KRASi group, yielding a set of seven genes (Figure 1C-D, S1E-G). Of these, the Axl receptor tyrosine kinase (*Axl*) showed the highest degree of enrichment, the lowest FDR = 0.002, and was found to be significantly positively enriched in 67% of all pairwise comparisons between KRASi and vehicle-treated tumors. AXL is a member of the TAM (TYRO3, AXL, and MERTK) family of receptor tyrosine kinases and has emerged as a key mediator of therapeutic resistance across multiple cancer types, including NSCLC(16) and PDAC (17). Upon activation by its ligand GAS6, AXL stimulates multiple downstream signaling networks, including the PI3K-AKT, JAK/STAT, and MAPK pathways, promoting cell survival and proliferation. Given the established role of AXL in therapeutic resistance, we investigated its potential as a co-target with KRAS inhibition. AXL protein was detected in all seven human NSCLC and seven murine KPP-derived LUAD cell lines examined (Figure 1E–F). In the KPP-derived LUAD cell line KP1, KRAS inhibition rapidly suppressed pERK and DUSP6 expression, with partial recovery at 24 hours and complete recovery by 72 hours (Figure 1G). This MAPK reactivation coincided with increased phosphorylation of AXL (Y779), implicating AXL signaling as a potential mediator of MAPK reactivation.

**Figure 1.**
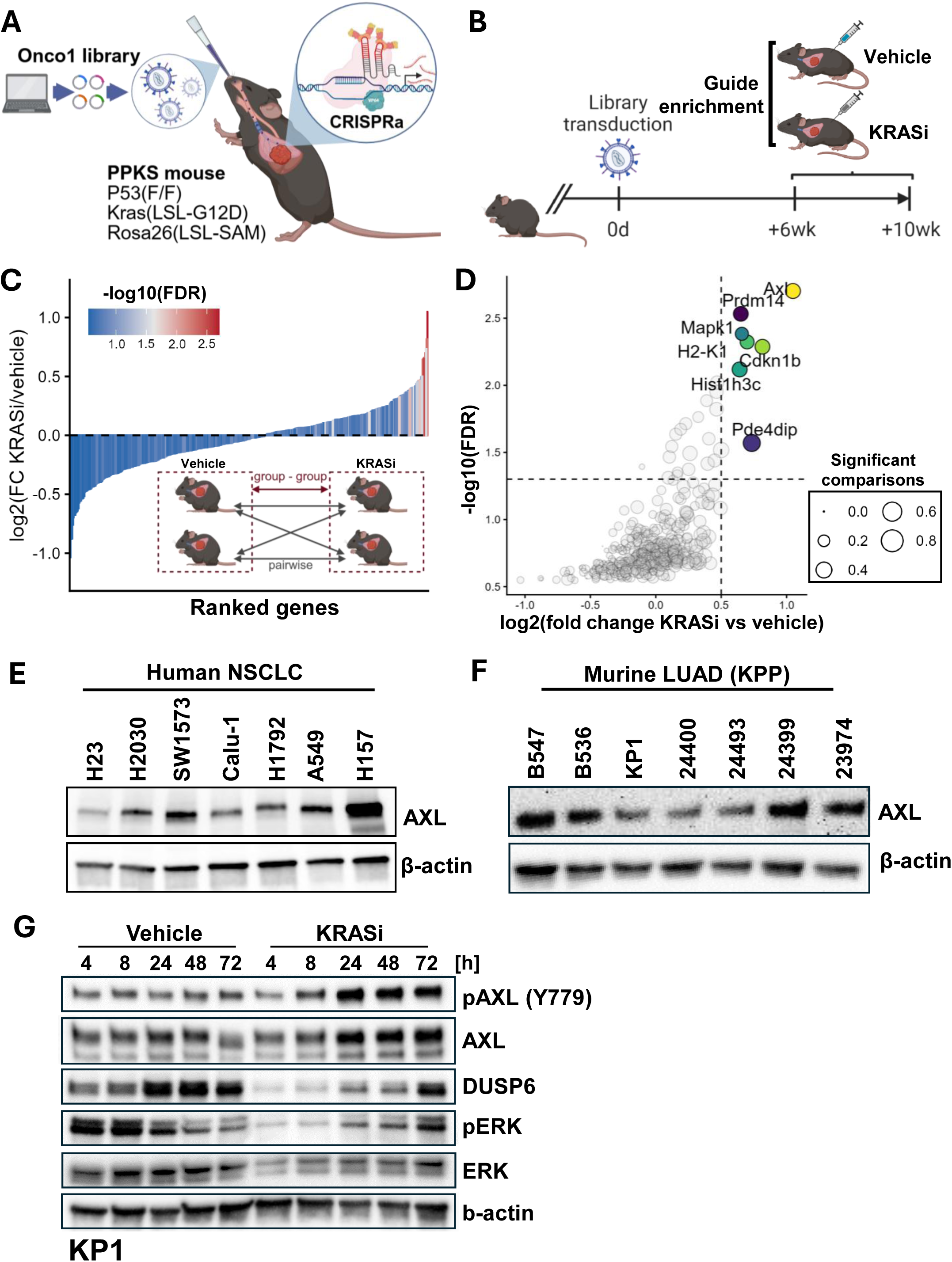
An unbiased in vivo CRISPR activation screen identifies AXL as a dominant driver of adaptive resistance to KRAS inhibition. A) Schematic of autochthonous in vivo CRISPRa screening. B) In vivo CRISPRa KRAS inhibitor screening scheme. C) Average gene-level enrichment in KRAS inhibitor treated tumors relative to vehicle controls. Inset schematic of group-to-group, and pairwise analysis. D) Significance and enrichment of genes in KRAS inhibitor-treated screening mice. Significant genes defined as log2(FC KRASi vs Vehicle) > 0.5 and FDR < 0.05. E) Immunoblot of AXL protein expression in seven human NSCLC cell lines. F) Immunoblot of AXL protein expression in seven KPP model-derived murine lung tumor cell lines. G) Immunoblot of AXL, phospho-AXL (pAXL), and MAPK-associated markers in response to KRAS inhibition (MRTX1133) or vehicle treatment in murine KP1 cells.

### Combined KRAS and AXL inhibition significantly deepens and extends MAPK suppression in KRAS mutant tumor cells

Several specific AXL inhibitors are currently undergoing clinical evaluation in NSCLC. Bemcentinib (R428/BGB324) is an orally available, selective inhibitor of AXL that binds to the intracellular catalytic domain of AXL and inhibits downstream signaling(18). To determine whether AXL inhibition enhances MAPK suppression, we treated cells with KRASi (MRTX1133) alone or combined with the AXL inhibitor bemcentinib (AXLi) and measured pERK, DUSP6, and MYC expression at 4 and 24 hours. Although MAPK suppression was similar at 4 hours, combination treatment resulted in consistently lower pERK, DUSP6, and MYC levels at 24 hours compared with KRASi alone (Figure 2A).

**Figure 2.**
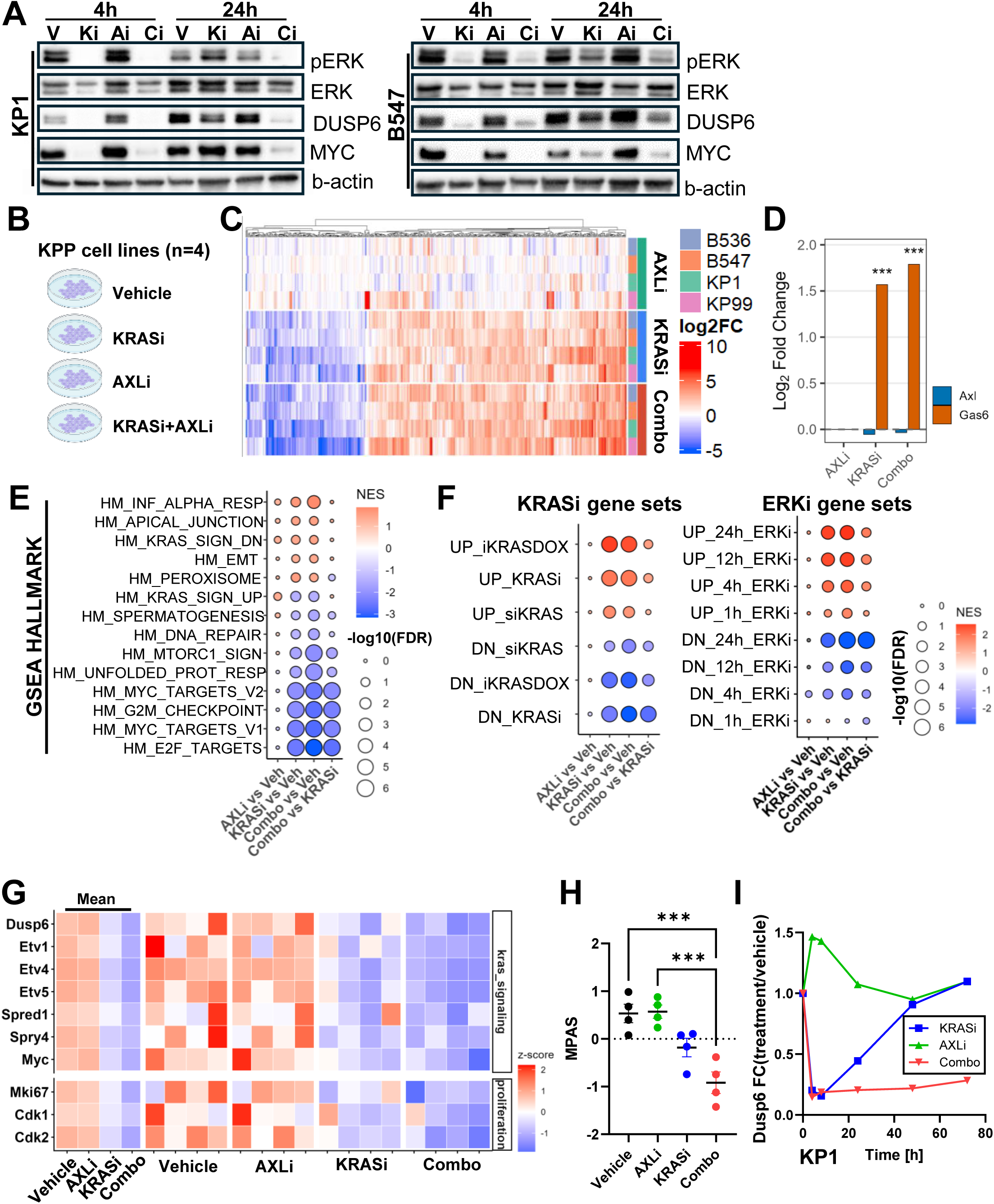
Combined KRAS and AXL inhibition deepens and extends MAPK suppression. A) Immunoblot of MAPK-markers in murine LUAD KP1 (left) and B546 (right) cells treated with KRASi (Ki; MRTX1133, 100nM), AXLi (Ai; bemcentinib, 500nM), alone or in combination (Ci), or with vehicle (V), harvested 4 or 24 hours after treatment initiation. B) Schematic of transcriptomic profiling of four murine LUAD (KPP) cell lines treated with KRASi, AXLi, alone or in combination, or with vehicle for 24 hours. C) Heatmap of all genes differentially expressed in combination-treated cells relative to paired vehicle controls. D) Differential expression of *Axl* and *Gas6* transcripts in treated cells. E) GSEA of significantly enriched HALLMARK gene sets in treated relative to vehicle controls. F) (left) GSEA of KRAS inhibition associated gene sets, (right) GSEA of ERK inhibition associated gene sets. G) Tile plot of KRAS and proliferation-associated genes in inhibitor-treated cells. H) MAPK Pathway Activity Score (MPAS) in KPP cells treated with KRAS and/or AXL inhibitor. Statistical testing performed using one-way ANOVA with multiple hypothesis correction. I) RT-qPCR time course of *Dusp6* mRNA in KP1 cells treated with KRASi, AXLi, alone or in combination, or with vehicle. *** indicates adjusted p < 0.001.

To characterize the transcriptional effects of combined KRAS and AXL inhibition, we performed mRNA-seq on four murine LUAD cell lines treated for 24 hours with KRASi, AXLi, KRASi/AXLi, or vehicle (16 samples total; Figure 2B). Although samples clustered primarily by cell line (Figure S2A), paired analysis relative to vehicle revealed extensive transcriptional changes following KRASi and combination treatment, with the greatest number of differentially expressed genes (DEGs) observed in the combination group (Figures 2C, S2B–C). While *Axl* expression was unchanged, *Gas6* was induced by both KRASi and combination treatment (Figure 2D). Most DEGs were shared between KRASi and combination treatment, suggesting that AXL inhibition largely augments the transcriptional response to KRAS inhibition. Consistent with this, 93.1% of shared DEGs exhibited a greater absolute fold change in the combination group than with KRASi alone (p = 2.79e-3; Figure S2D). Notably, genes showing the largest differential response to combination treatment included canonical MAPK targets such as *Etv1*, *Etv4*, and *Etv5*, supporting enhanced suppression of KRAS/MAPK signaling (Figure S2E).

To investigate this effect further, we conducted gene set enrichment analysis (GSEA) and gene set variation analysis (GSVA) using the HALLMARK gene sets, which revealed enhanced suppression of multiple KRAS and proliferation-associated gene sets in combination-treated cells (Figure 2E and S2F).

We also analyzed a set of recently described KRAS and ERK gene sets that capture the effect of MAPK inhibition with high fidelity(18,19), which revealed enhanced enrichment in combination-treated samples (Figure 2F and S2G). Overall, the GSEA and GSVA results were consistent with the addition of AXL inhibition enhancing the transcriptomic effect of KRAS inhibition.

To further investigate the effect of combination treatment on KRAS transcriptomic activity, we examined the expression of six KRAS-associated genes (including *Etv4*, *Dusp6*, and *Myc*), revealing profound suppression in combination-treated cells (Figure 2G), which was corroborated with qRT-PCR (Figure S2H-I). We also noted significantly suppressed expression of proliferation markers (*Mki67* and *Cdk1*) in combination-treated samples relative to KRAS inhibition alone. To quantify transcriptomic MAPK activity as a function of treatment, we calculated the clinically relevant MAPK Pathway Activity Score (MPAS) for each sample(20), which revealed significantly lower MPAS in the combination group relative to KRASi alone (Figure 2H).

To assess the durability of pathway suppression, we measured *Dusp6* and *Myc* expression following treatment. While both genes returned to baseline within 48 hours with KRASi alone, suppression persisted for at least 72 hours with KRASi/AXLi combination treatment (Figures 2I and S2J). Because *Gas6* expression increased in response to treatment, suggesting compensatory autocrine signaling, we examined whether the GAS6-AXL axis is associated with acquired KRASi resistance. Reanalysis of three public RNA-seq datasets from human NSCLC (H23, H358) and PDAC (Panc0203, PANC-1) models with acquired resistance to sotorasib, ARS-1620, or MRTX1133 revealed upregulation of both AXL and GAS6, with GAS6 showing the strongest and most uniform induction across cell lines and inhibitors (Figure S3). In summary, these results show that the combination of KRAS and AXL inhibition results in enhanced and sustained suppression of MAPK signaling and that GAS6 overexpression may play a role in MAPK reactivation.

### KRASi and AXLi synergistically suppress tumor cell viability *in vitro*

To test the therapeutic potential of combined KRASi and AXLi *in vitro*, we quantified the baseline response of six human NSCLC cell lines to daraxonrasib (RMC-6236), a KRAS MULTI(ON) tri-complex inhibitor with proven clinical efficacy in NSCLC and PDAC, and to AXL inhibitor (Figure 3A and S4A). To assess synergy between KRAS and AXL inhibition, we performed combinatorial treatment and synergy assays, which revealed synergy between RMC-6236 and AXLi in a majority of cell lines (Figure 3B-C, S4B-C), and strong synergy between adagrasib and AXLi in 2 of 4 KRAS^G12C^ cell lines (Figure S4D-E). We also found synergy between KRASi (MRTX1133) and AXLi in the human KRAS^G12D^ PDAC cell line HPAF-II (Figure 3D). We assessed the effect of combined KRASi (adagrasib) and AXLi on clonogenic potential in the NSCLC cell line H23, which revealed enhanced suppression (Figure 3E). To validate these findings in cross-species analysis, we profiled nine murine LUAD (KPP) and nine murine PDAC (KPC) tumor-derived cell lines. This revealed widely varying responses to KRASi, while the responses to AXLi were more uniform (Figure 3F and Figure S4F). Synergy analysis revealed significant synergistic interactions across a range of drug concentrations in 17 of 18 cell lines (Figure 3G-H and S4G-J). We also found that combinatorial treatment significantly reduced the clonogenic potential relative to KRASi alone across a set of seven murine LUAD cell line, (Figure 3I and S5A). We next quantified the effect of drug treatment using live-cell imaging and endpoint staining in three independent murine LUAD cell lines (KP1, 24399, and B536). These experiments revealed proliferation defects in combination-treated cells (Figure S5B-C), with significantly reduced proliferation (Figure 3J). We noted that the presence of abnormal nuclei (abnormal shape and multi-nucleation) was significantly higher in combination-treated wells (Figure S5D), and we found that combination-treated nuclei were significantly enlarged (Figure 3K). To determine if sensitivity to the combination was retained in cell lines with acquired resistance to KRASi, we tested three murine LUAD cell lines and four murine PDAC cell lines with matching resistant subclones (Figure 3L and S5E). We found that the synergistic effect of combined treatment was retained in 2 of 3 resistant lung, and 3 of 4 resistant pancreatic subclones (Figure 3M and Figure S5F-G). Notably, both, resistant lung subclones that showed retained synergy also displayed higher levels of phospho-AXL than their naïve counterparts (Figure 3N).

**Figure 3.**
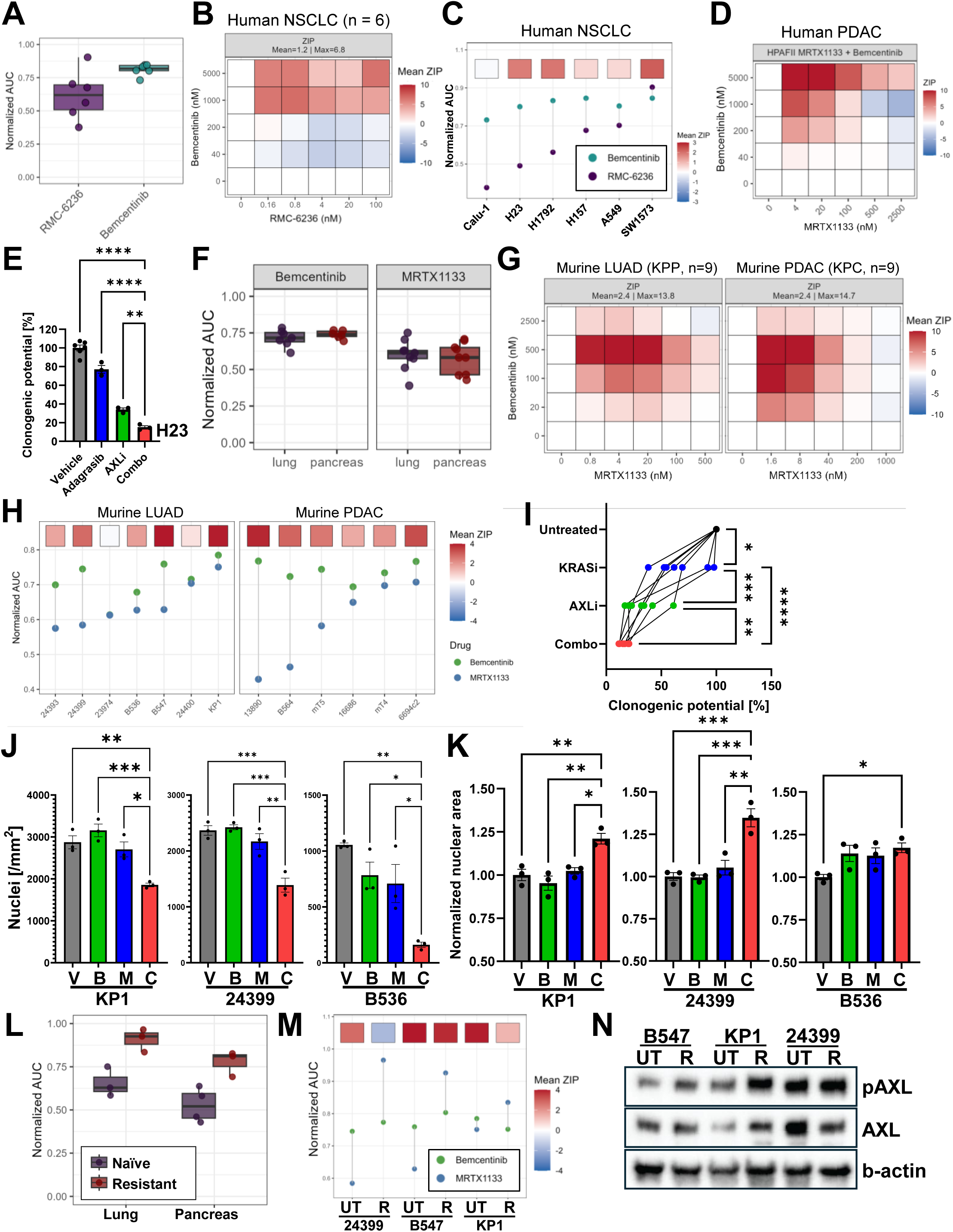
KRAS and AXL inhibition synergize across KRAS-targeted therapies and tumor models. A) Normalized AUC for KRAS inhibitor (daraxonrasib; RMC-6236) and AXL inhibitor (bemcentinib) in a panel of six human NSCLC cell lines, B) Ensemble average ZIP synergy scores for daraxonrasib and AXLi in six human NSCLC cell lines, C) Normalized AUC and mean ZIP score in NSCLC cell lines, D) ZIP-synergy analysis of KRASi (MRTX1133) and AXLi in the human pancreatic cancer cell line HPAF-II, E) Clonogenic assay of KRASi (adagrasib) and AXLi in human H23 NSCLC cells, F) Normalized AUC for KRAS inhibitor (MRX1133) and AXLi in a panel of murine LUAD (KPP) and PDAC (KPC) cell lines. G) Ensemble average ZIP synergy scores for KRASi (MRTX1133) and AXLi across 9 murine LUAD (left) and 9 murine PDAC (right) cell lines. H) Normalized AUC and mean ZIP score in murine LUAD (left) and PDAC (right) panel. I) Clonogenic potential assay in KRASi/AXLi-treated murine LUAD cell lines (n = 7), J) Number of nuclei per mm2 after 48 hours of KRASi/AXLi-treatment, K) normalized nuclear area after 48 hours of KRASi/AXLi-treatment. L) Normalized AUC for KRASi (MRTX1133) in murine LUAD and PDAC cells with (resistant) and without (naïve) acquired resistance to KRASi. M) Normalized AUC and mean ZIP score for KRASi (MRTX1133) and AXLi-treated murine LUAD cells with and without acquired resistance, N) Immunoblot of pAXL and AXL in murine LUAD cells with and without acquired resistance to KRASi. (B, D and G) Absolute synergy scores capped at 10 for visualization purposes.

### Combined KRAS and AXL inhibition reduces tumor growth and extends MAPK suppression in multiple syngeneic KRAS mutant models

To assess the potential of combined KRAS and AXL inhibition in vivo, we evaluated treatment responses in several syngeneic subcutaneous transplantation models (Figure 4A). First, we used the murine LUAD lung tumor model-derived B536 cell line, a KRASi and AXLi-responsive cell line that displays an intermediate level of synergy with AXL inhibition (Figure 3H). We implanted 500,000 B536 cells bilaterally into the flanks of C57BL/6 mice and when the average tumor size reached 80-100 mm^3^, the mice were randomized to receive KRASi (MRTX1133, 10 mg/kg, i.p., q.d.) and/or AXLi (bemcentinib, 50 mg/kg, p.o., b.i.d.), or vehicle control. While KRASi significantly reduced tumor growth, the effect was enhanced in combination-treated tumors, with regression observed in 12 of 16 tumors and reduced tumor weights at treatment endpoint (Figure 4B-D). To test a more challenging scenario, we repeated the experiment with the murine LUAD cell line KP1, which exhibits the highest levels of intrinsic KRASi and AXLi resistance in our murine cell line panel (Figure 3H). In addition to the MRTX1133 treated cohort, we also treated mice with RMC-236 (6 mg/kg, p.o. q.d.) alone or in combination with AXLi to simulate likely clinical translation. We found that while KRAS inhibition exerted a significant tumor-restrictive effect, the addition of AXLi increased the effect of both MRTX1133 and RMC-6236 and resulted in significantly reduced tumor volumes and weights (Figure 4E and 4F). Consistent with a high degree of intrinsic resistance, we did not observe tumor regression in this model. We found that combination-treated KP1 tumors were significantly less proliferative than KRASi-treated tumors, as indicated by reduced fraction of Ki67-positive cells (Figure 4G-H). To assess MAPK-inhibition in vivo, we performed IHC for phospho-ERK1/2 (pERK) on tissues harvested 4 hours after the last treatment (Figure 4G and 4I). We found that KRAS inhibitor-treated tumors showed no loss of phospho-ERK1/2; in fact, phosho-ERK1/2 staining was enhanced in these samples, indicating compensatory ERK activation and potential development of resistance. Remarkably, a significant fraction (6/10) of combination-treated samples showed minimal pERK staining, consistent with sustained MAPK suppression. 4/10 combination-treated tumors showed pERK staining comparable to that of vehicle control tumors, indicating heterogeneity in response. We also performed mRNA-seq on these subcutaneous tumors (5 per group, 20 tumors in total, see Figure S6A-C). These results revealed transcriptomic suppression of KRAS/MAPK-associated genes and gene sets and a comparable reduction in MPAS in KRASi and combination-treated tumors, indicative of discrepancy between pERK and transcriptomic MAPK activity (Figure S6D-G). These results show that combined KRASi and AXLi potently suppresses tumor growth across models and inhibitor classes.

**Figure 4.**
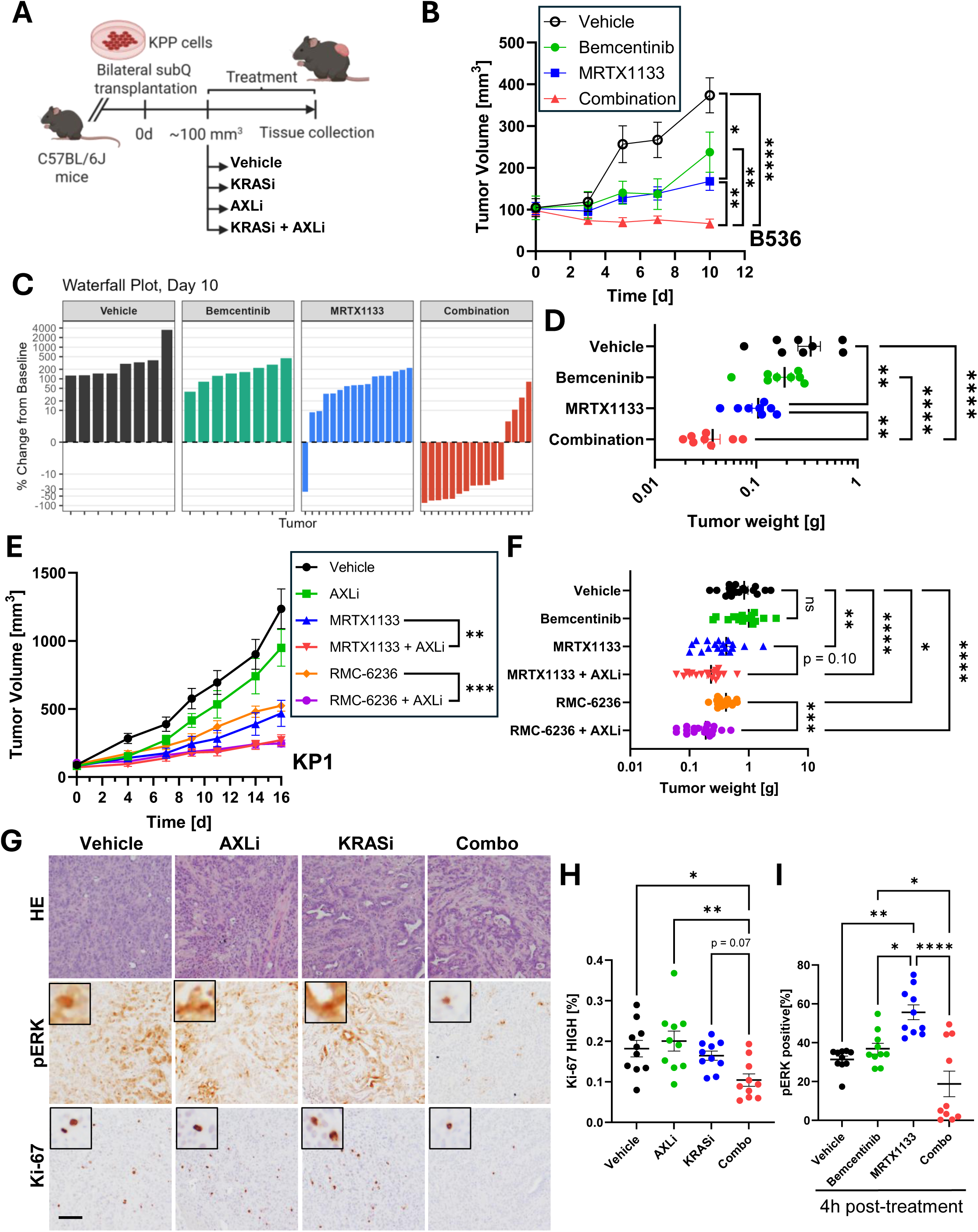
Combined KRAS and AXL inhibition enhances tumor control in syngeneic KRAS-driven lung cancer models. A) Schematic of subcutaneous syngeneic murine LUAD (KPP) tumor transplantation KRASi (MRTX1133 or RMC-6236) and AXLi (bemcentinib) treatment response model. B) Subcutaneous B536 tumor volume as function of time in mice treated with KRASi (MRTX1133), AXLi, alone or in combination, or with vehicle. C) Tumor volume waterfall plot in B536 subcutaneous model. D) Tumor weight was function of treatment in the B536 model. D) Tumor weights from B536 model. E) Subcutaneous KP1 tumor volume as function of time in mice treated with KRASi (MRTX1133 or RMC-6236), AXLi, alone or in combination, or with vehicle. F) Tumor weights from B536 model. G) Histology and IHC staining (pERK and Ki-67) in KP1 tumors harvested 4 hours after the final treatment. KRASi; MRTX1133, scale bar 100 um, H) Quantification of proliferation (Ki-67) in subcutaneous KP1 tumors. I) Quantification of pERK in subcutaneous KP1 tumors. (B, E) tumor volumes were analyzed using a linear mixed-effects model with fixed effects and p values were adjusted for multiple testing using the Benjamini-Hochberg method, (D, F, H, I) Statistical testing performed using one-way ANOVA with multiple hypothesis correction, * p<0.05, ** p<0.01, *** p<0.001, **** p<0.0001.

### Combined treatment drives an interferon gamma response in vivo

To further characterize the response to therapy, we next performed GSEA of the Hallmark gene sets on the KRASi and AXLi-treated KP1 tumors, which revealed strong enrichment for several interferon and immune reaction-related gene sets in all treatment groups relative to vehicle controls, with enhanced enrichment in combination-treated tumors relative to KRASi alone (Figure 5A). We also found that a set of IFNγ-related transcripts were significantly over-expressed in combination-treated tumors (including *Ifng*, *Ifngr1*, *Stat1*, and several MHC class I genes, see Figure 5B). In addition we noted that transcripts encoding the T-cell recruiting chemokines *Cxcl9* and *Cxcl10*, and various T-cell markers (such as *Cd3*e and *Cd8a*) were significantly overexpressed in both KRASi and combination-treated tumors, with combination-treated tumors showing higher fold changes (Figure 5C). To further investigate treatment-induced change in the tumor microenvironment, we conducted scRNA-seq using 10X Genomics FLEX chemistry from pooled FFPE samples from each treatment group (8 pooled samples in total). Following sample integration and clustering, we identified a large epithelial/tumor cell cluster, as well as clusters representing cancer-associated fibroblasts (CAFs), tumor-associated macrophages/dendritic cells, and a relatively small cluster of combined T cell/NK cells (Figure S7A-C). We found that combination-treated cells in the tumor cell cluster displayed over-expression of a large set of IFNγ-induced genes, including canonical IFNγ signaling members (such as *Stat1* and *Irf1*), antigen-processing factors (such as *Tap1* and *Tap2*), MHC class I genes, chemokines, and inflammatory markers (Figure 5D). We calculated an IFNγ module score using the expression of 15 IFNγ-inducible genes, revealing increased scores in epithelial cells from combination-treated tumors (Figure 5E and S7D). These findings were corroborated using pseudobulk analysis of the tumor cell cluster (Figure S7E-F). We also observed increased expression of IFNγ-regulated genes in the TAM/DC cluster (Figure 5F). Unfortunately, the T/NK-cell cluster contained too few cells to subcluster further (likely due to technical issues with FFPE sample processing for FLEX chemistry). However, we found that the expression of T cell markers (including *Cd3g* and *Cd8a*) increased in combination-treated samples while NK cell-associated markers decreased (Figure 5G-H).

**Figure 5.**
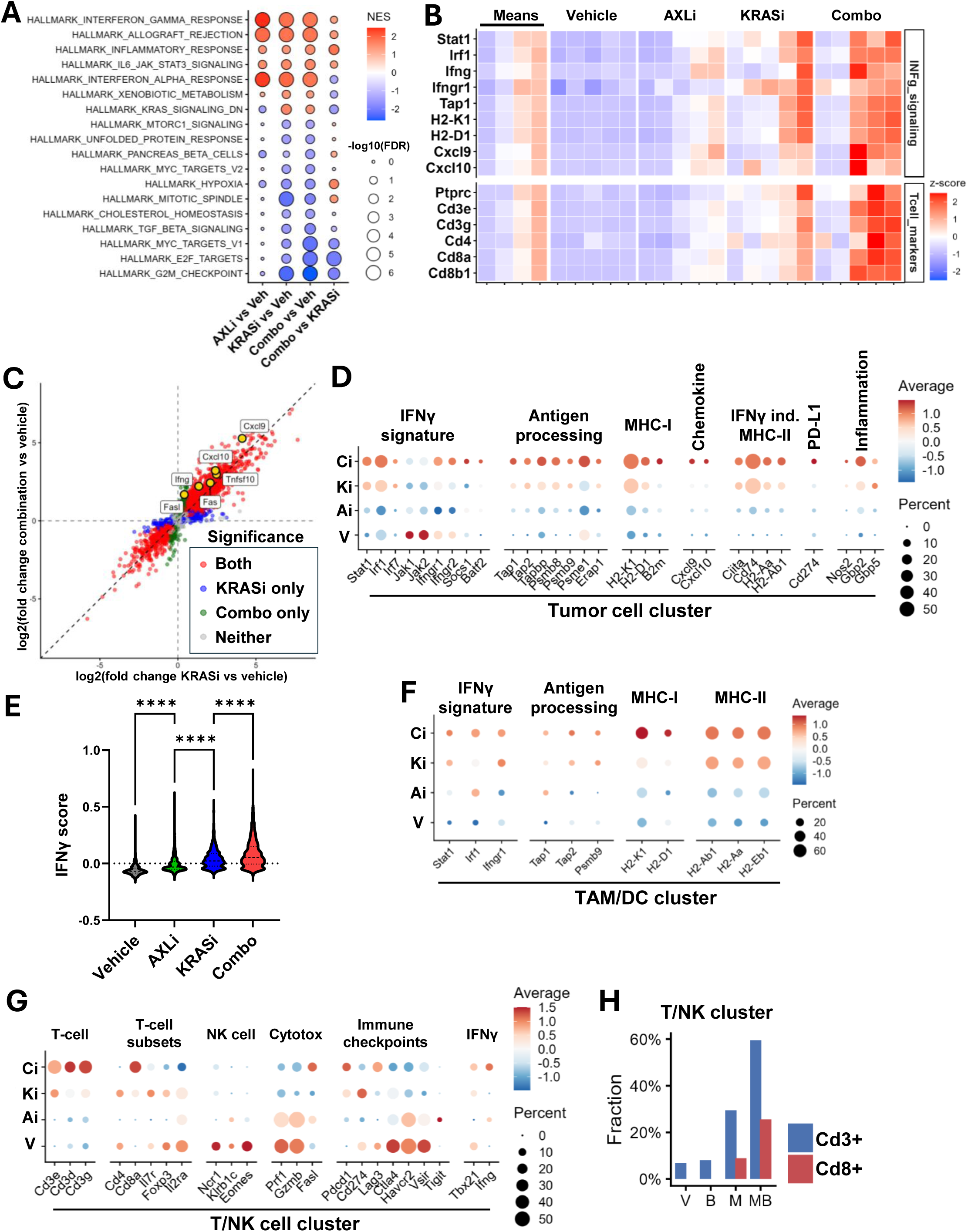
AXL inhibition amplifies KRAS inhibitor-induced inflammatory and immune responses. A) GSEA of significantly enriched Hallmark gene sets in subcutaneous KP1 tumors treated with KRASi (MRTX1133), AXLi, alone or in combination, or with vehicle. B) Tile plot of bulk RNA-seq expression of IFNγ-regulated and T-cell marker genes. C) Expression fold-change vs vehicle for combination and KRAS inhibitor treated tumors with a set of combination-enhanced IFNγ-induced genes highlighted in yellow. D) Dot plot of IFNγ and immune associated gene expression in the epithelial/tumor cell cluster from scRNA-seq. E) IFNγ regulated gene expression score in tumor cell cluster. F) Dot plot of IFNγ associated gene expression in the TAM/DC cell cluster. G) Dot plot of T-cell associated gene expression in the T-cell/NK-cell cluster. H) Quantification of fraction Cd3 and Cd8 expressing cells in the T-cell/NK-cell cluster.

### Combined KRAS and AXL inhibition enhances recruitment of cytotoxic T cells to the tumor microenvironment

We next conducted quantitative IHC staining, with the intention of orthogonal validation of scRNA seq analyses as well as to overcome some of the artifacts associated with loss of immune cells contexture with FLEX chemistry. The analysis revealed increased phospo-Stat1 (pStat1) and MHC class I staining (both commonly induced in response to IFNγ) in combination-treated tumors (Figure 6A-C and S8A). We also found that combination-treated tumors contained significantly more CD45+, and CD8a+ cells (Figure 6A and 6D-F). To independently validate these findings, we implanted a second cohort of mice with KP1 cells and performed spectral flow cytometry using a 21-marker immune cell-profiling panel on tumors harvested after short-term treatment (6 days, see Figure 6G and Figure S8B-C). This confirmed that combination-treated tumors contain more CD8+ T cells (Figure 6H). Within the CD8+ T cell subsets, we also found that combination-treated tumors were enriched for effector memory CD8+ T cells (Figure 6I). Both KRASi and combination-treated tumors displayed increased CD86 expression in cDC1 cells and a shift in polarization toward an M1-like (pro-inflammatory) macrophage phenotype (Figure 6J-K), consistent with enhanced antigen presentation and a tumor-restrictive microenvironment.

**Figure 6.**
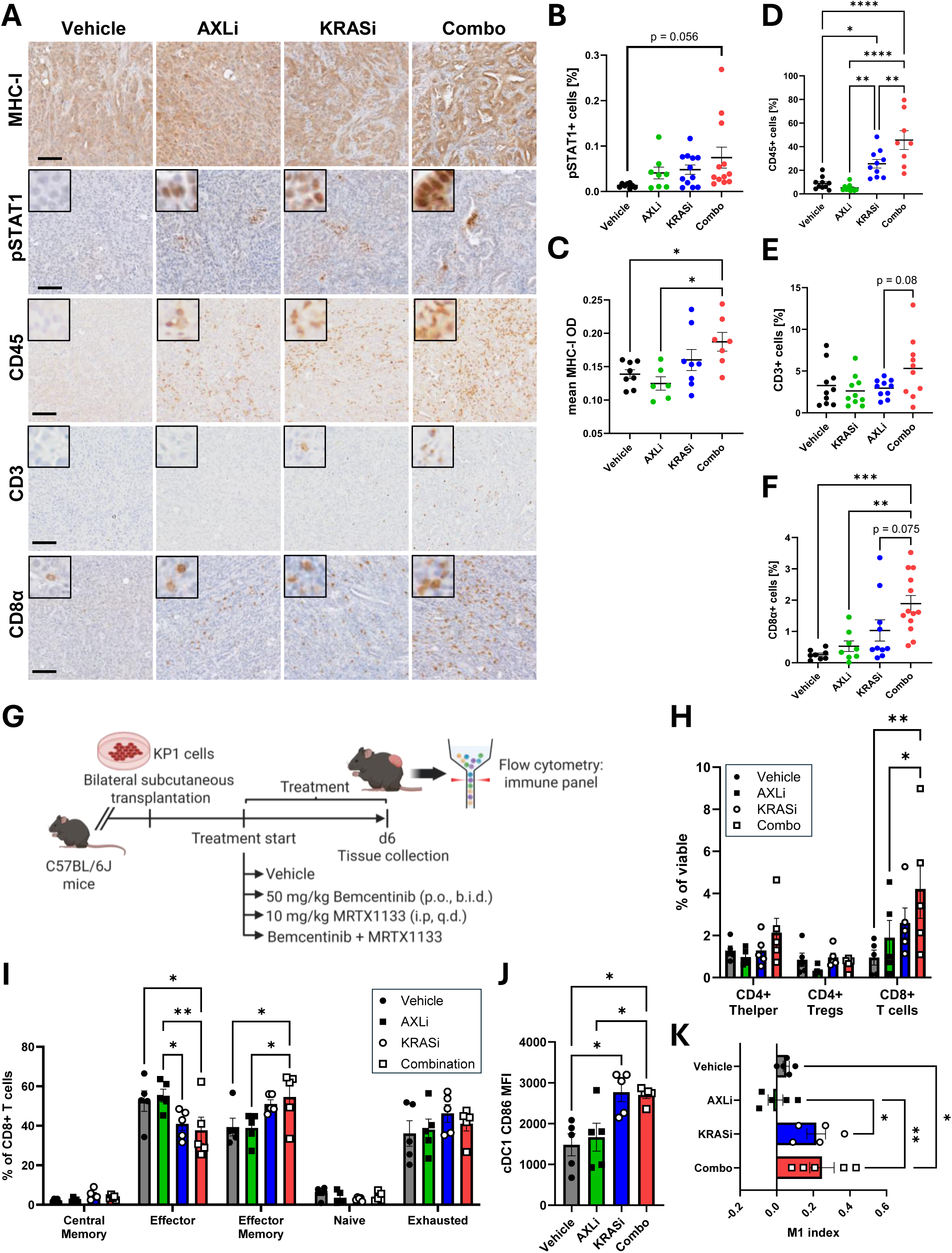
KRAS and AXL co-targeting drives tumor immune remodeling and T-cell recruitment. A) IHC staining of MHC class I, pSTAT1, CD45, CD3 and CD8a in subcutaneous KP1 KRASi (MRTX1133)/AXLi-treated tumors, scale bar 100 um. B) Quantification of pSTAT1+ cell fraction [%] in subcutaneous tumors. C) Quantification of MHC-I staining (optical density; OD) in tumor samples. D) Quantification of CD45+ cell fraction [%] in subcutaneous tumors. E) Quantification of CD3+ cell fraction [%] in subcutaneous tumors. F) Quantification of CD8a+ cell fraction [%] in subcutaneous tumors. G) Schematic of immune profiling timepoint KP1 tumor treatment response experiment. H) T cell subset abundance as function of treatment. I) CD8 T cell subset abundances. J) cDC1 cell CD86 staining intensity (median fluorescent intensity; MFI). K) M1 macrophage index; (M1-M2)/(M1+M2). Statistical testing performed using one-way ANOVA (B-F, J, K) or two-way ANOVA (H, I) with multiple hypothesis correction, * p<0.05, ** p<0.01, *** p<0.001, **** p<0.0001.

### KRASi/AXLi-treatment and IFNγ synergistically induce FAS-mediated tumor cell killing

While our in vitro results show that combined KRASi and AXLi have a synergistic tumor-cell autonomous effect on cell viability and proliferation, the activation of IFNγ signaling and enhanced recruitment of T-cells in vivo indicate that tumor immunity likely also plays a role in the response in vivo. T cells and NK cells are the primary sources of IFNγ in the tumor microenvironment(21) and exert anti-tumor activity through FAS-mediated tumor cell killing, among other mechanisms. Furthermore, FAS overexpression and FAS-mediated tumor control have been previously reported in response to KRAS inhibition [PMID: 37625401]. Consistent with this, we found that combination-treated tumors displayed overexpression of *Fas* and *Fasl*, and decreased expression of *Cflar* (encoding for c-FLIP, an antagonist of FAS-mediated apoptosis) (Figure 7A-B). Interestingly, we found that combination-treatment significantly increases *Fas* and decreases *Cflar* expression relative to KRASi alone *in vitro* (Figure 7C). In fact, *Fas* expression displayed one of the highest deltas upon comparison of combinations versus KRASi fold-changes (Figure S2E). Furthermore, we found that combination treatment significantly increased FAS protein expression and surface presentation of FAS (Figure 7D-E). Importantly, we did not detect *Ifng* transcript expression in any of our KPP cells in vitro, indicating that tumor-cell autonomous FAS overexpression in response to treatment is not directly driven by autocrine IFNγ signaling. However, the type II interferon receptor subunits *Ifngr1* and *Ifngr2* were both overexpressed in combination-treated cells, suggesting enhanced sensitivity to exogenous IFNγ stimulation (Figure 7C). Given the strong induction of IFNγ signaling in the combination-treated tumors and the potential of enhanced IFNγ response in combination-treated cells, we next assessed the effect of IFNγ on drug-treated KPP cells. We found that both KP1 and B536 strongly upregulated MHC class I in response to IFNγ and that the response was enhanced in combination-treated cells (Figure 7F and S9A-C). While IFNγ alone had a modest effect on FAS, the combination of KRASi/AXLi and IFNγ dramatically increased surface presentation of FAS protein in both cell lines, suggesting that KRASi/AXLi/IFNγ-treated cells may be poised to respond strongly to FAS-stimulation (Figure 7G-I and S9D-F). To assess if IFNγ renders KRASi/AXLi-treated cells more susceptible to FAS-mediated apoptosis, we conducted live-cell imaging experiments using a fluorogenic label responsive to activated effector Caspases 3 and 7 and stimulated the cells with an agonistic FAS antibody. These results showed minimal induction of apoptosis in the absence of FAS-stimulation or IFNγ, but remarkably strong induction of apoptosis in KRASi/AXLi/IFNγ/anti-FAS-treated cells (Figure 7J-K and S9G).

**Figure 7.**
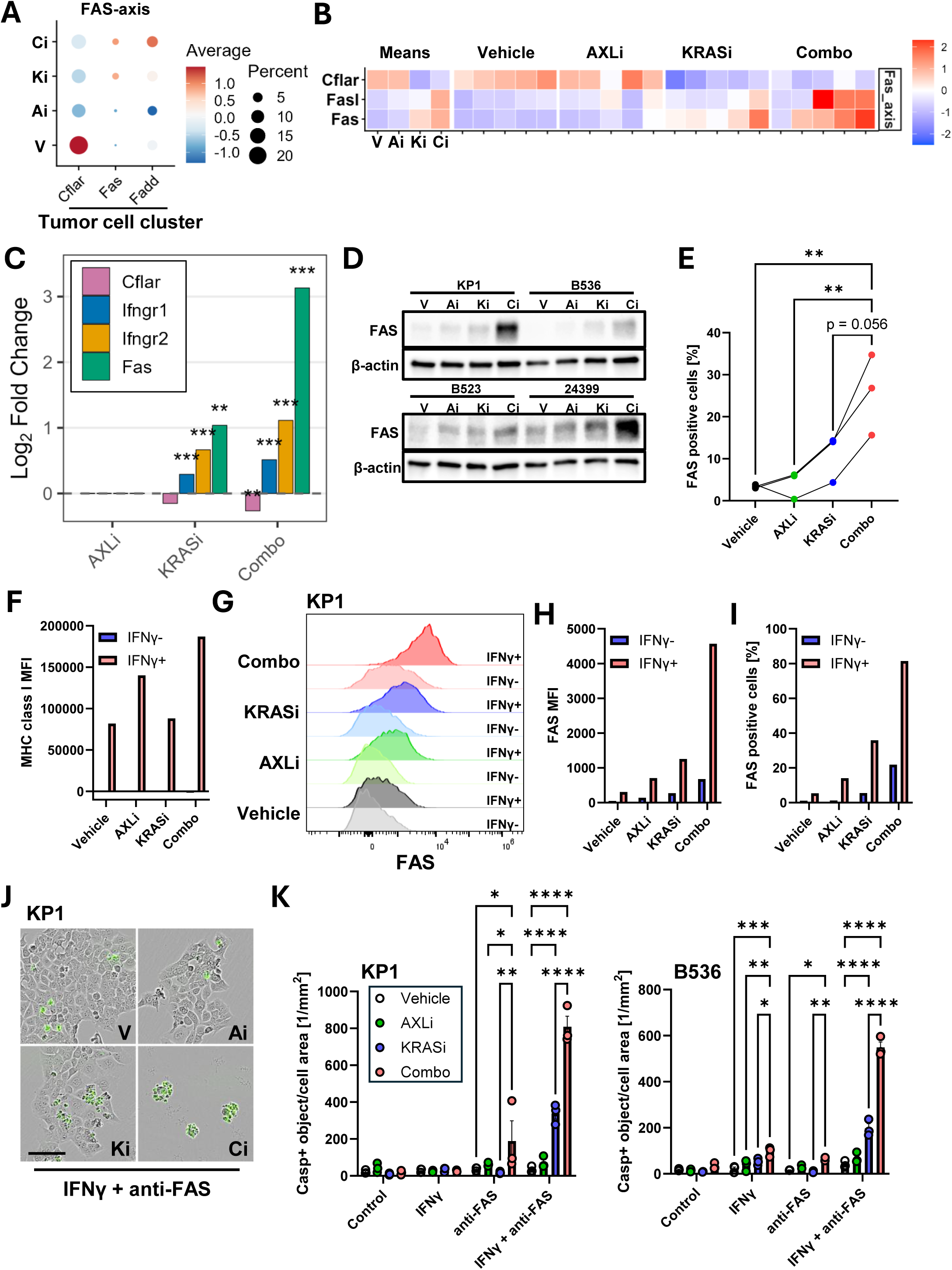
Combined KRAS and AXL inhibition sensitizes tumor cells to IFNγ- and FAS-mediated apoptosis. A) Dot plot of FAS-axis genes in KP1 tumor cell cluster from scRNA-seq, B) Tile-plot of FAS-axis genes from KP1 tumor bulk RNA-seq, C) Differential expression of FAS-axis (Fas, Cflar) and IFNã receptor transcripts in KRASi/AXLi-treated KPP cells in vitro, D) Immunoblot of FAS protein in KRASi/AXLi-treated murine LUAD cells, E) Fraction FAS-positive murine LUAD cells as function of KRASi and/or AXLi treatment assessed with flow cytometry. F) MHC class I surface presentation (median fluorescent intensity; MFI) as function of treatment in KP1 cells. G) Flow cytometry histograms of FAS surface protein presentation in KP1 cells treated with KRASi/AXLi, with and without IFNγ, H) FAS surface presentation (median fluorescent intensity; MFI) as function of treatment in KP1 cells. I) Fraction FAS+ cells as function of treatment, J) Fluorescence microscopy of KP1 cells treated with KRASi/AXLi, IFNγ and agonistic FAS antibody, fluorogenic Caspase activity label in green. Scale bar 100 um.

These results provide a mechanistic framework showing that combining KRAS with AXL inhibition suppresses tumor growth through both tumor cell autonomous and immune modulatory mechanisms, defining a promising avenue to extend therapy response.

## Discussion

Direct KRAS inhibitors have transformed the treatment of KRAS-mutant cancers, but adaptive resistance remains a major limitation. Although several RTKs have been implicated in this process, it has remained unclear whether resistance reflects numerous context-specific bypass pathways or converges on a smaller set of dominant regulators in vivo. Using an unbiased CRISPRa screen in autochthonous KRAS-driven lung tumors, we identified AXL as a dominant RTK mediator of adaptation to KRAS inhibition, whereas other receptors previously linked to resistance, including EGFR, ERBB2, FGFR1, MET, and PDGFRA, were not similarly enriched. While these receptors may remain important in specific contexts, our findings suggest that adaptation in the native tumor environment converges on a more limited set of signaling nodes and highlight the value of studying therapeutic resistance directly in vivo.

AXL inhibition alone had little effect on MAPK signaling or tumor-cell proliferation, consistent with AXL being largely dispensable while oncogenic KRAS remains active. In contrast, the role of the AXL/GAS6 axis became apparent only after KRAS blockade. Initial ERK suppression was similar with KRAS inhibition alone and with the combination, but dual treatment significantly delayed the subsequent recovery of phospho-ERK, DUSP6, and MYC. Rather than deepening the immediate pharmacodynamic response, AXL inhibition delayed the rebound that followed. This distinction argues against AXL acting as a constitutive driver in these models and instead defines it as a conditional vulnerability created by therapy. Notably, an independent study by Ching et al. likewise identified AXL as a mediator of resistance to RAS pathway inhibition in KRAS-mutant pancreatic and lung cancer models and demonstrated enhanced antitumor activity with combined RAS and AXL inhibition, providing orthogonal validation of AXL as a broadly relevant adaptive node following KRAS pathway suppression(22).

Furthermore, these molecular effects translated across experimental systems, including cross species studies. Combined KRAS and AXL inhibition suppressed proliferation and clonogenic growth in human NSCLC and PDAC cell lines, as well as in multiple independently derived murine lung and pancreas cancer cell lines and remained active in KRAS inhibitor-resistant subclones with elevated phospho-AXL. The interaction was also not confined to one KRAS allele or inhibitor class. AXL inhibition enhanced responses to RAS(ON) inhibitor daraxonrasib/RMC-6236 in human and murine cell lines, to MRTX1133 in KRAS^G12D^ murine and human pancreatic cancer cells, and to adagrasib in KRAS^G12C^ lung cancer cells. Combined KRAS and AXL inhibition not only prolonged MAPK suppression but also amplified IFNγ-responsive and antigen-presentation programs, increased CXCL9/CXCL10 expression, and promoted cytotoxic T-cell infiltration. These findings suggest that AXL coordinates both tumor-cell-intrinsic adaptation and suppression of antitumor immunity, positioning it at the intersection of the signaling and immune responses to KRAS inhibition. While a companion by Ching et al. similarly found that AXL inhibition enhanced the efficacy of KRAS pathway blockade, our data further suggests that immune remodeling contributes to this response(22).

The FAS/FASL axis may help explain these effects. Consistent with prior studies showing that KRAS inhibition sensitizes tumors to FAS-dependent T-cell killing(15), combined treatment increased *Fas* expression, elevated *Fasl*, and reduced the anti-apoptotic regulator *Cflar*. *Fas* induction occurred even in cultured tumor cells lacking IFNγ stimulation, indicating a tumor-intrinsic priming for death-receptor signaling. Moreover, IFNγ rendered combination-treated cells highly sensitive to FAS activation *in vitro*. *In vivo*, this increased susceptibility coincided with enhanced antigen presentation and greater CD8+ T-cell infiltration. Together, these findings suggest that dual KRAS/AXL inhibition both sensitizes tumor cells to immune attack and promotes a microenvironment capable of executing it, providing a plausible mechanism for improved tumor control.

AXLi improved responses to several KRAS-inhibitors with distinct allele selectivity and mechanisms of action, arguing that AXL dependence reflects a common response to pathway suppression. As broader RAS inhibitors enter clinical development, targeting a shared adaptive regulator may prove more useful than building combinations around each individual KRAS allele. The existing clinical experience with AXL inhibitors also lowers an important barrier to translation. Biomarker development will nevertheless be essential, particularly because basal AXL abundance may be less informative than treatment-induced AXL and GAS6 activation or the capacity for MAPK rebound.

However, several limitations remain. The CRISPR activation library was enriched for recurrently altered cancer genes and therefore did not capture the full range of epigenetic, metabolic, and non-coding mechanisms that may support adaptation. Additional work in patient-derived models and clinical specimens will be needed to establish how frequently tumors become AXL-dependent and which features predict benefit from the combination. Moreover, phospho-ERK was not durably suppressed in every tumor, indicating that other escape routes remain active and may eventually limit combined KRAS and AXL inhibition. Defining those secondary mechanisms will be important for understanding both primary nonresponse and acquired resistance.

AXL emerges from this work as a central mediator of adaptation to KRAS inhibition. By promoting MAPK pathway recovery while restraining antitumor immunity, AXL enables tumors to persist under therapeutic pressure. Disrupting this induced dependency prolonged pathway suppression, enhanced immune activation, and improved efficacy across multiple KRAS-targeted therapies. These findings identify AXL as a clinically actionable combination partner and support evaluation of KRAS/AXL co-inhibition to improve the durability of responses in KRAS-mutant cancers.

### Methods Mouse models

All studies were conducted in compliance with institutional guidelines (MD Anderson; IACUC). PPKS (*p53^F/F^;Kras^LSL-G12D/+^;Rosa26^LSL-SAM/+^*) mice were generated and housed at MD Anderson. C57BL/6 mice for subcutaneous implantation were purchased from The Jackson Laboratory. All mice received standard chow diet *ad libitum* and were housed in pathogen-free facility with standard controlled conditions. No more than 5 mice were housed together, all mice were under the supervision of veterinarians in an AALAC-accredited animal facility at the University of Texas M.D. Anderson Cancer Center. All animal procedures were reviewed and approved by the MDACC IACUC (ACUF #00001626, PI: Maitra).

### In vivo CRISPRa screening

In vivo CRISPRa screening was performed largely as described previously, with the addition of KRAS inhibition (cite CRM). Thirteen PPKS mice were infected by nasal instillation with pooled sgOnco1/CMV-Cre lentivirus. After 6 weeks, mice were randomized to receive MRTX1133 (3 mg/kg daily, intraperitoneally) or vehicle (10% SBE-β-CD, 50 mM citrate buffer, pH = 5) for 4 weeks before tissue collection. Genomic DNA was isolated from lungs (DNeasy Blood and Tissue Kit, Qiagen), and guide sequences were amplified by PCR with dual-indexed primers to generate NGS libraries, which were sequenced on a NextSeq2000 (Illumina). Guide counts were extracted from FASTQ files using a modified published script (PMID: 28333914), normalized, and analyzed with DESeq2 to calculate average fold changes (KRASi vs. vehicle). Gene-level enrichment was assessed using CRISPhieRmix (PMID: 30296940). Genes were considered significant if they met all of the following criteria: log2 fold change > 0.5, FDR < 0.01 based on averaged counts, and significant enrichment (FDR < 0.1) in at least 50% of pairwise KRASi-versus-vehicle comparisons.

### KPP and KPC model-derived murine cell lines

Murine lung tumor cell lines were derived from KPP (Kras^LSL-G12D^, P53^F/F^), KP (Kras^LSL-G12D^, P53^F/wt^) or KPPY (Kras^LSL-G12D^, P53^F/F^, Rosa26^LSL-YFP^) tumor-bearing mice treated with adenoCre or lentiCre through nasal instillation. KPC cell lines were derived from KPC (Kras^LSL-G12D^, P53^F/wt^, Ptf1a^Cre^) or KPCY (Kras^LSL-G12D^, P53^F/wt^, Ptf1a^Cre^, Rosa26^LSL-YFP^) tumor-bearing mice. The PKCY-derived 6694c2 cell line was purchased from Kerafast. The KPC-derived cell lines mT4 and mT5 were kind gifts from David Tuveson (CSHL).

### Cell Culture

Murine tumor cell lines and the human PDAC (HPAF-II) and NSCLC (SW1573, Calu-1, A549, H23, H157 and H1792) cell lines were maintained and propagated in DMEM/10% FBS/1% Penicillin-Streptomycin.

### KRAS inhibitor resistant cell lines

MRTX1133 cell lines were generated by treating cells with increasing concentrations of drug over an extended period of time. Treatment was initiated at the estimated IC50 value for each cell line and gradually increased at each passage until the cells proliferated unhindered in the presence of 1 uM MRTX1133.

### Immunoblot analysis

Protein lysates were collected in RIPA lysis buffer with HALT protease and phosphatase inhibitor at indicated timepoints. Mouse and human tumor cells were processed for immunoblotting using a previously described protocol (Thege et al 2022). For immunoblotting analysis, the cultures were treated with vehicle (DMSO), 100 nM MRTX1133, 500 nM bemcentinib or combination 24 hours after seeding. Antibodies are listed in Supplementary Table and blots were analyzed using a ChemiDoc XRS (BioRad).

### mRNA-seq of inhibitor treated cell lines

Murine KPP tumor cells (KP1, KP99, B536, and B547) were treated with MRTX1133 (100 nM), bemcentinib (500 nM), their combination, or DMSO for 24 hours. Total RNA was isolated (RNeasy Mini Kit, Qiagen), and poly(A)-selected libraries were sequenced (150 bp paired-end) on an Illumina NovaSeq X Plus (Novogene). Transcript abundance was quantified from FASTQ files using Salmon against the mm10 transcriptome. Differential expression analysis was performed with DESeq2 following filtering of lowly expressed protein-coding genes (total count >10). A paired design (∼ cell_line + group) was used to account for cell line effects. Differential expression was assessed using Wald tests for AXLi vs vehicle, KRASi vs vehicle, combination vs vehicle, and combination vs KRASi comparisons, with adjusted *p* < 0.05 considered significant. Log2 fold changes were shrunk using the *ashr* method. Genes with baseMean >10 were retained for downstream analyses. Significant genes from the combination-versus-vehicle comparison were used for heatmap generation from log2-transformed normalized counts. Ranked log2 fold-change values were used for GSEA (v4.2.2), while variance-stabilized data were used for GSVA. KRASi and ERKi gene sets were obtained from previous studies (PMIDs: 38843329, 38843331), and MAPK Pathway Activity Scores were calculated as described previously (PMID: 29872725).

### Analysis of deposited RNA-seq datasets

RNA-seq count data from GSE229070, GSE164326 and GSE269985 were analyzed in R using DESeq2. Genes were annotated using the supplied references and filtered to retain only protein-coding genes. Samples were matched to metadata, and lowly expressed genes (total counts ≤10 across all samples) were excluded. Differential expression analysis was performed using a DESeq2 model that included cell line and treatment group effects (∼ cell_line + group). Log2 fold-change estimates for the resistant-versus-vehicle comparison were shrunk using the apeglm method.

### RT-qPCR

RNA from mouse tumor lines treated with MRTX1133 and/or bemcentinib, or vehicle, were harvested in RLT buffer with beta mercaptoethanol and isolated using the RNeasy Mini kit (Qiagen) with on-column DNAse treatment following the manufacturer’s instructions at indicated time points. cDNA was generated using BioRad iScript kit, and qPCR was performed using Power SYBR Green PCR Master Mix on a QuantStudio 3 thermocycler (Thermo Scientific). Primer sequences are listed in Supplementary Table. Relative gene expression was calculated using the ddCt method with β-actin as housekeeping gene. When indicated, gene expression was normalized to vehicle-treated control.

### Quantification of single compound and synergistic drug response

Drug response was assessed as previously described (Thege et al., 2022). Cells were seeded in 96-well plates (10,000 human or 2,500 murine cells/well) and treated with varying concentrations of KRAS inhibitors (RMC-6236, MRTX1133, or adagrasib) and/or bemcentinib for 72 hours. Cell viability was measured by MTT assay, with absorbance read at 570 nm (690 nm reference). Single-agent dose-response curves were fitted using a four-parameter log-logistic model (LL.4). From these fits normalized area under the curve (AUC) was calculated by integrating predicted viability over the log-transformed dose range and scaling to the maximal response window. Matrix synergy scores were calculated using normalized viability values with SynergyFinder in R returning ZIP, Bliss, Loewe and HSA scores. Mean ZIP scores across concentrations and cell lines were calculated.

### Clonogenic assay

The clonogenic potential of both mouse and human tumor lines was determined utilizing a standard 12-well clonogenic assay. Cells were seeded at 1000/well and allowed to adhere for 24 hours before the addition of media containing varying concentrations of inhibitor. After 10 days of incubation with drug, plates were stained with crystal violet. Plates were imaged after staining using a ChemiDoc XRS (BioRad), and the colony area was analyzed using ImageJ software.

### IncuCyte proliferation assay

Cell proliferation was assessed using an IncuCyte live-cell imaging system, with proliferation estimated based on confluency measurements. Cells were seeded at a density of 2,500 cells per well in 100 µL of media. The following day, drug treatments (DMSO vehicle, MRTX1133, bemcentinib, or combination) were added and plates were imaged every 2 hours, capturing four fields per well, over a period of 4 days. **End-point proliferation assay and assessment of nuclear features**

Cell proliferation and morphology were assessed using an endpoint imaging assay. Cells were seeded at a density of 2,500 cells per well in 100 µL of media and allowed to adhere overnight. After 48 hours of incubation with drug, media were removed and cells were fixed with 4% paraformaldehyde. Cells were permeabilized using 0.1% Triton X-100, 1% BSA, and 40 µg/mL RNase, then stained with propidium iodide, phalloidin–iFluor488 in 1% BSA/PBS. Plates were imaged using an IncuCyte imaging system. Nuclear number and features (nuclear area) were quantified using CellProfiler (v.4.2.8). Abnormal nuclei (abnormal shape, multi-nucleation and nuclear fragmentation) were manually counted.

### IncuCyte FAS-mediated apoptosis assay

Cells were seeded as described above. Following over-night incubation, the media was replaced with complete media containing DMSO vehicle, MRTX1133 (10nM), bemcentinib (500nM), MRTX1133 + bemcentinib, IFNγ (5ng/ml), and/or anti-FAS/proteinG (anti-FAS; 5ug/ml, protein G; 2ug/ml), as indicated, in addition to CellEvent Caspase-3/7 Green (1 uM). The plates were imaged every 4 hours, capturing four fields per well, over a period of 48 hours. The number of Caspase positive objects was normalized to the total cell-containing area in each image.

### Syngeneic subcutaneous tumor model

For assessment of tumor response, C57BL/6J mice were implanted bilaterally in the flanks with 500,000 B536 or KP1 cells suspended in a 1:1 mix of Matrigel and PBS. Tumors were measured with calipers every 2-3 days and tumor volumes estimated as V = L*W^2^/2. When the average tumor volume reached 80-100 mm^3^, the mice were randomized to receive MRTX1133 (10 mg/kg, i.p. q.d., in 50 mM citrate buffer pH 5, 10% SBE-β-CD and 0.9% NaCl), RMC-6236 (6 mg/kg, p.o. q.d. in 0.5% HPMC, 0.1% Tween-80 in water), bemcentinib (50 mg/kg p.o. b.i.d. in 0.5% HPMC, 0.1% Tween-80 in water), MRTX1133 and bemcentinib, RMC-6236 and bemcentinib, or vehicle as indicated. The tissues were harvested 4 hours after the last treatment, fixed in 10% neutral buffered formalin, flash frozen and/or used for flow cytometry.

### Tumor growth analysis

Tumor volume data were analyzed in R using the lme4, lmerTest, and emmeans packages. Tumor volumes were natural log-transformed and analyzed using a linear mixed-effects model with fixed effects. The significance of model terms was assessed by analysis of variance. Post hoc pairwise comparisons between treatment groups at each time point were performed using estimated marginal means, with p values adjusted for multiple testing using the Benjamini-Hochberg method.

### Quantitative IHC and histology

IHC and hematoxylin and eosin (H&E) staining were performed as previously described (Thege et al 2022). Tissue sections (5 µm) were stained with H&E or used for IHC using standard histologic methods. IHC was performed using citrate (pH = 6.0) or Tris-EDTA antigen retrieval. Antibodies used are listed in Supplementary table. Quantification of staining areas was done using Aperio ImageScope (v.12.4.3.5008) and Qupath (v.0.5.1). For assessment of MHC class I upregulation, all identified cell objects were exported from Qupath and analyzed further in R.

### mRNA-seq of subcutaneous tumors

Total RNA was isolated from flash-frozen subcutaneous tumors using the RNeasy Mini Kit (Qiagen) with on-column DNase treatment. Poly(A)-selected mRNA libraries were sequenced (150 bp paired-end) on an Illumina NovaSeq X Plus (Admera). Reads were aligned with STAR, and differential expression (treatment vs. vehicle) was analyzed using DESeq2. Ranked log2 fold-change values were used for GSEA (v4.2.2; GenePattern, Broad Institute), and MAPK Pathway Activity Scores were calculated as previously described.

### scRNA-seq of subcutaneous tumors

Single-cell RNA-seq was performed on 1.5-mm FFPE mouse lung tissue biopsies using the Chromium GEM-X Flex Gene Expression 16-plex platform (10x Genomics). Following tissue processing and dissociation, viability-adjusted cell suspensions were pooled and used for GEM generation, barcoding, and library preparation. Libraries were sequenced on an Illumina NovaSeq X Plus to a mean depth of ∼25,000 reads per cell (Novogene). Data were processed with Cell Ranger v9.0 using the GRCm39 reference transcriptome. Quality control, decontamination (DecontX), doublet removal (scDblFinder), normalization (SCTransform), integration, clustering, and cell-type annotation were performed using Seurat. Cell-type subsets were reclustered following normalization. Pseudobulk differential expression analysis of epithelial/tumor cells was performed by aggregating raw counts at the sample level using Seurat’s AggregateExpression function and analyzing filtered count matrices with DESeq2 using default settings.

### Tumor cell preparation for flow cytometry

Subcutaneous tumors (two per mouse) were collected and bisected. Half of each tumor was pooled for flow cytometry, while the remaining tissue was fixed in 10% neutral-buffered formalin or flash-frozen. Tumors were mechanically dissociated and digested in DMEM containing collagenase IV (1 mg/mL) and DNase I (50 U/mL) for 35 min at 37°C with agitation. Cell suspensions were filtered through 100-µm strainers, washed, and resuspended in FACS buffer (1% BSA, 2 mM EDTA in PBS) for staining and analysis.

### Spectral flow cytometry

Single cell suspensions were first stained with a fixable viability dye 700 (BD) and Fc receptor blocking antibody (anti-CD16/CD32, Biolegend) for 20 min at 4 °C in the dark. Following washing, cells were incubated with surface antibodies targeting CD45, CD4, CD8, CD90.2, CD44, CD62L, PD-1, TIM-3, LAG-3, CD39, CD103, CD11b, CD11c, MHC class I/II, CD86, and CX3CR1, for 30 min at 4 °C. Subsequently, cells were fixed and permeabilized using Foxp3 fixation/permeabilization buffer (Invitrogen) and incubated with intracellular antibodies targeting Foxp3, Ki-67, Granzyme B, and TOX. For Fas/MHC class I experiments, cells were stained with FAS and MHC class I antibodies. Fluorescence minus one (FMO) controls were included where appropriate for low-intensity markers to ensure accurate gating strategies. Flow cytometry data was acquired on a Cytek Aurora 5L cytometer and analyzed by FlowJo Software (v10.10.1).

### Data Availability

The sequencing data generated in this study are publicly available in NCBI GEO at GSE307283, GSE307369 and GSE307368. Additional datasets were obtained from NCBI GEO at GSE269985, GSE229070 and GSE164326.

## Acknowledgements

F.I.T. is supported by a Seed Grant from the Hirshberg Foundation for Pancreatic Cancer Research. S.M.W. is supported by the German Cancer Aid, a Max-Eder Junior Group Leader Award (70114858), a Department of Defense, Pancreatic Cancer Research Program Idea Development Award (HT94252310855) and the European Research Council (ERC) Starting Grant (ADRIP-ERC-2024-STG) under the European Union’s Horizon Europe research and innovation programme (GAP101163784). A.M. is supported by grants from the NIH (TBEL; U54CA274371, PASSCODE; U24CA274274, and EDRN; U01CA200468). The MDACC Flow Cytometry and Cellular Imaging (FCCI) Core Facility, Small Animal Imaging Facility (SAIF), and DVMS Veterinary Pathology Core are supported by a Cancer Center Support Grant (P30CA016672) from the NIH. This research was supported by The Ohio State University Comprehensive Cancer Center (OSUCCC) under a Cancer Center Support Grant (P30CA016058) from the NIH. This research was also made possible through support provided by the Pelotonia Institute for Immuno-Oncology (PIIO), which is funded by the Pelotonia community and the OSUCCC.

---

**Figure S1.**
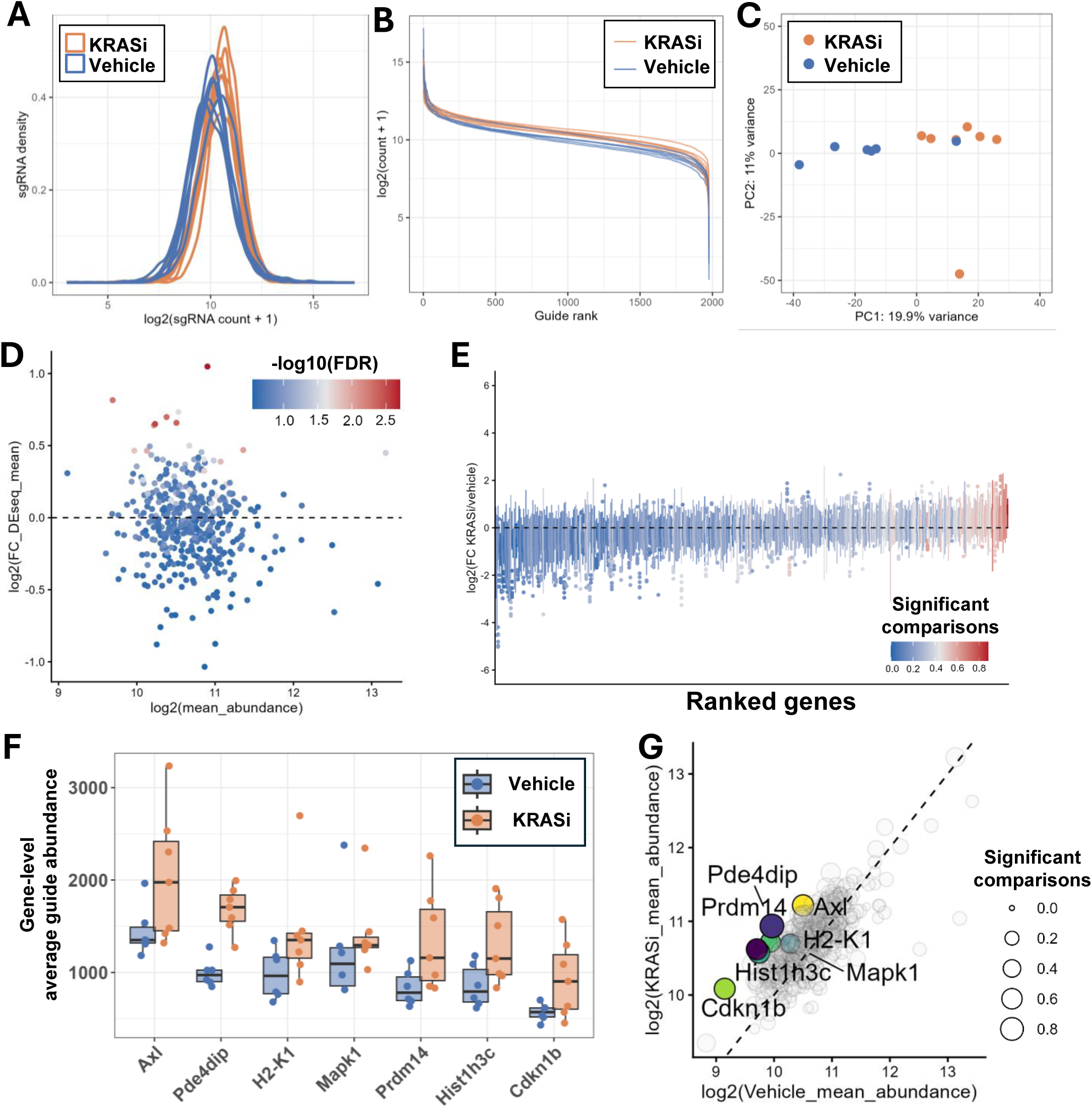
Validation of the autochthonous in vivo CRISPR activation screen. Supplemental for. Figure 1. A) Guide distribution in KRAS inhibitor and vehicle-treated Onco1 screening tumor samples. B) Guide abundance (log2(normalized count +1)) vs guide rank. C) guide-level Principal Component Analysis. D) MA-plot (log2(fold-change) vs log2(mean guide abundance)). E) Per-tumor gene-level fold-change enrichment in KRAS inhibitor treated tumors relative to vehicle controls F) Gene-level abundance of the significant genes identified in the screen. G) KRAS inhibitor sample vs vehicle sample abundance with significant genes highlighted.

**Figure S2.**
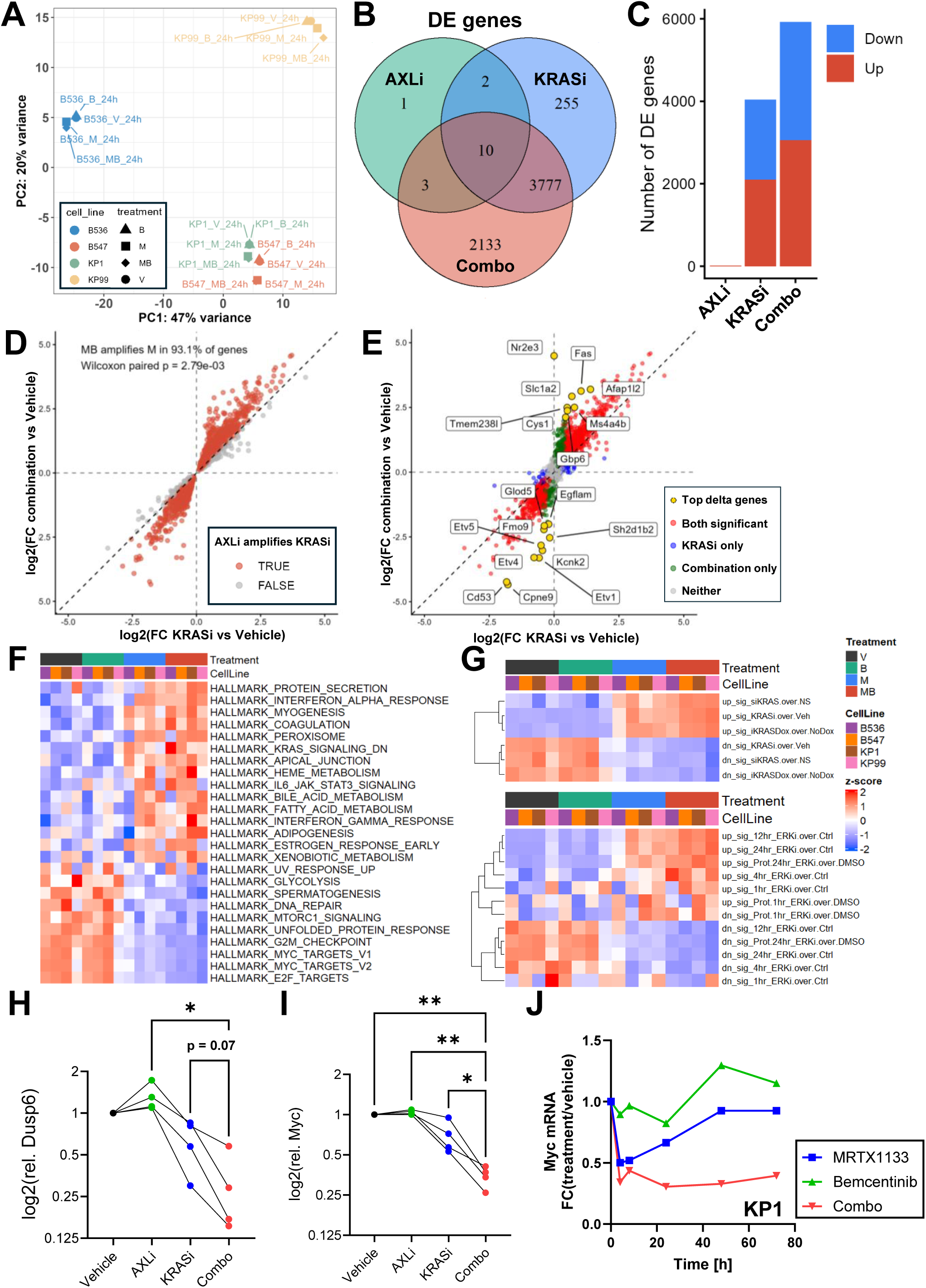
Combined KRAS and AXL inhibition prevents adaptive MAPK reactivation in tumor cells. Supplemental for. Figure 2. A) PCA plot for mRNA-seq of KRASi (MRTX1133) and/or AXLi-treated murine LUAD (KPP) cell lines. B) Venn diagram of differentially expressed genes (pairwise, relative to vehicle controls). C) Number of differentially expressed genes relative to vehicle for each treatment group. D) Fold-change vs vehicle for combination and KRAS inhibitor treated samples. Genes where combination treatment enhanced differential expression over KRAS inhibitor alone highlighted in red. E) Fold-change vs vehicle for combination and KRAS inhibitor treated samples. The 20 genes with the highest combination vs KRASi delta (log2FC(Combination) - log2FC(KRASi)) highlighted in yellow. F) GSVA results for Hallmark gene sets in KRASi/AXLi-treated murine LUAD cells. G) GSVA results for KRASi (top) and ERKi (bottom) associated gene sets. H) RT-qPCR analysis of Dusp6 mRNA in murine LUAD cells treated with KRASi, AXLi, alone or in combination, or with vehicle, harvested 24 hours after treatment initiation. I) RT-qPCR analysis of Myc mRNA in murine LUAD cells. RT-qPCR time course of Myc mRNA in KP1 cells as function of treatment. Statistical testing performed with one-way ANOVA, * p < 0.05, ** p < 0.01.

**Figure S3.**
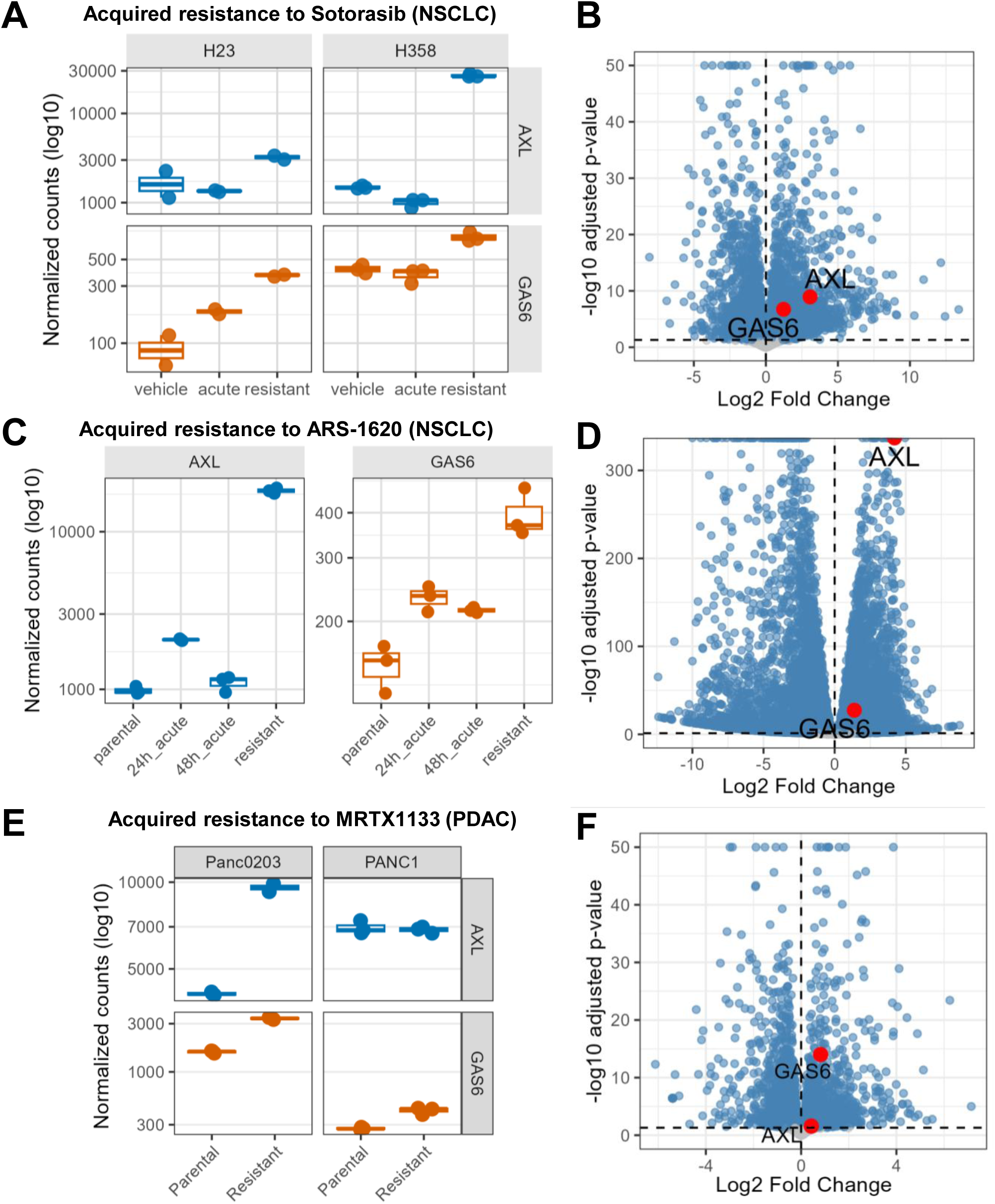
AXL and GAS6 are induced during acquired resistance to KRAS inhibition. Supplemental for. Figure 2. A) AXL and GAS6 transcript expression (log10(normalized counts)) in human NSCLC cells lines H23 and H358 in response to acute KRAS inhibition with sotorasib, or with acquired resistance. B) Volcano plot highlighting AXL and GAS6 in resistant vs vehicle-treated cells. C) AXL and GAS6 expression H358 cells in response to acute KRAS inhibition with ARS-1620, or with acquired resistance. D) Volcano plot highlighting AXL and GAS6 in resistant vs vehicle-treated cells. E) AXL and GAS6 expression human PDAC Panc0203 and PANC1 cells with acquired resistance to MRTX1133, and naïve controls. F) Volcano plot highlighting AXL and GAS6 in resistant vs vehicle-treated cells. (A-B) data from GSE229070, (C-D) data from GSE164326, (E-F) data from GSE269985.

**Figure S4.**
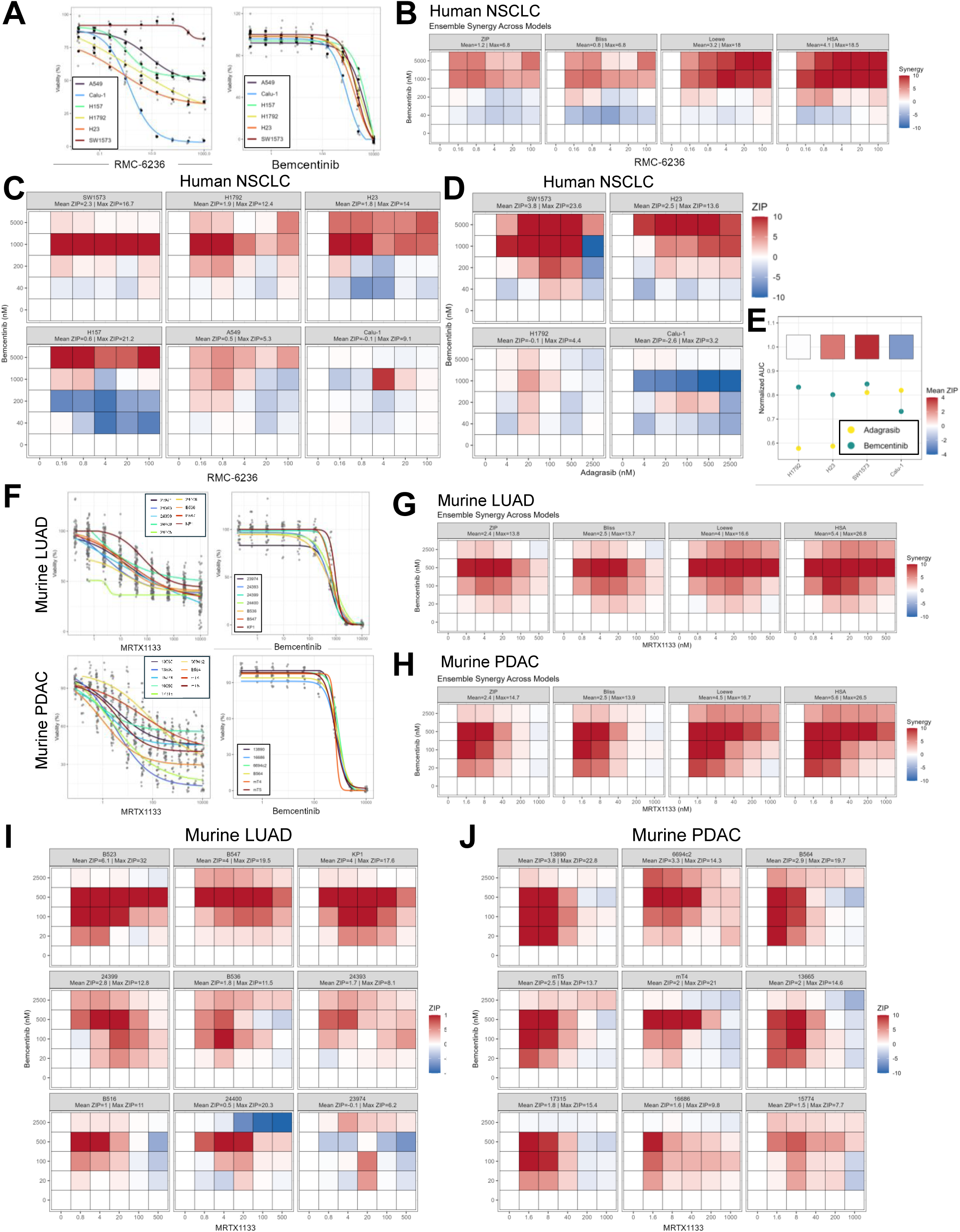
Synergistic responses to KRAS and AXL co-targeting across tumor models. Supplemental for. Figure 3. A) Drug response curves for KRASi (RMC-6236, left) and AXLi (right) in a panel of six human NSCLC cell lines, B) Ensemble average synergy scores (ZIP, Bliss, Loewe and HSA) for RMC-6236 and AXLi in human NSCLC cell lines, C) RMC-6236/AXLi ZIP synergy scores for individual human NSCLC cell lines, D) Adagrasib/AXLi ZIP synergy scores for individual KRASG12C mutant human NSCLC cell lines, E) Normalized adagrasib and AXLi AUC and mean ZIP synergy scores in KRASG12C mutant human NSCLC cell lines, F) Drug response curves for KRASi (MRTX1133) and AXLi in murine LUAD and PDAC cell lines, G) Ensemble average synergy scores for KRASi (MRTX1133) in nine murine LUAD cell lines, H) Ensemble average synergy scores for KRASi (MRTX1133) in nine murine PDAC cell lines, I) KRASi (MRTX1133)/AXLi ZIP synergy scores for individual murine LUAD cell lines, J) KRASi (MRTX1133)/AXLi ZIP synergy scores for individual murine PDAC cell lines. (B-D and G-J) Absolute synergy scores capped at 10 for visualization purposes.

**Figure S5.**
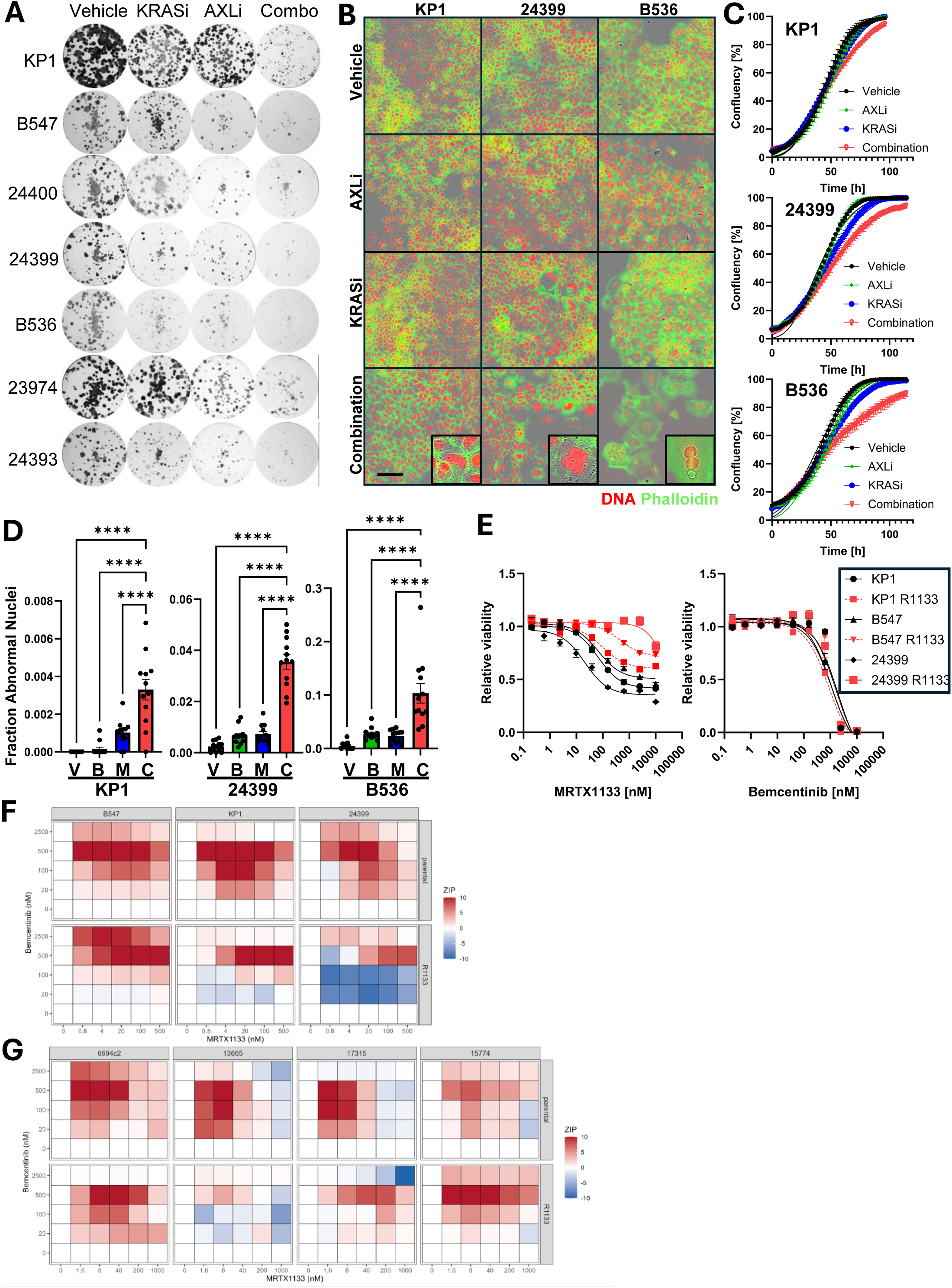
Combined KRAS and AXL inhibition enhances antitumor activity in vitro. Supplemental for. Figure 3. A) Clonogenic assay in seven murine LUAD (KPP) cell lines treated with KRASi (MRTX1133) and/or AXLi, or vehicle. B) Fluorescent staining of KPP cells treated with inhibitors for 48 hours, (red – propidium iodide, green – phalloidin), scale bar 100 um. C) Confluency as function of time in response to KRASi, AXLi and combination treatment for three KPP cell lines. D) Fraction of abnormal nuclei as function of treatment, 48 hours after treatment initiation. E) Dose response curves for KRASi (MRTX1133, left) and AXLi (right) in KPP cell with and without acquired resistance (R1133) to MRTX1133. F) KRASi (MRTX1133)/AXLi ZIP synergy scores of paired naïve (top) and MRTX1133 resistant (bottom) murine LUAD cell lines. F) KRASi (MRTX1133)/AXLi ZIP synergy scores of paired naïve (top) and MRTX1133 resistant (bottom) murine PDAC cell lines. (F-G) Absolute synergy scores capped at 10 for visualization purposes.

**Figure S6.**
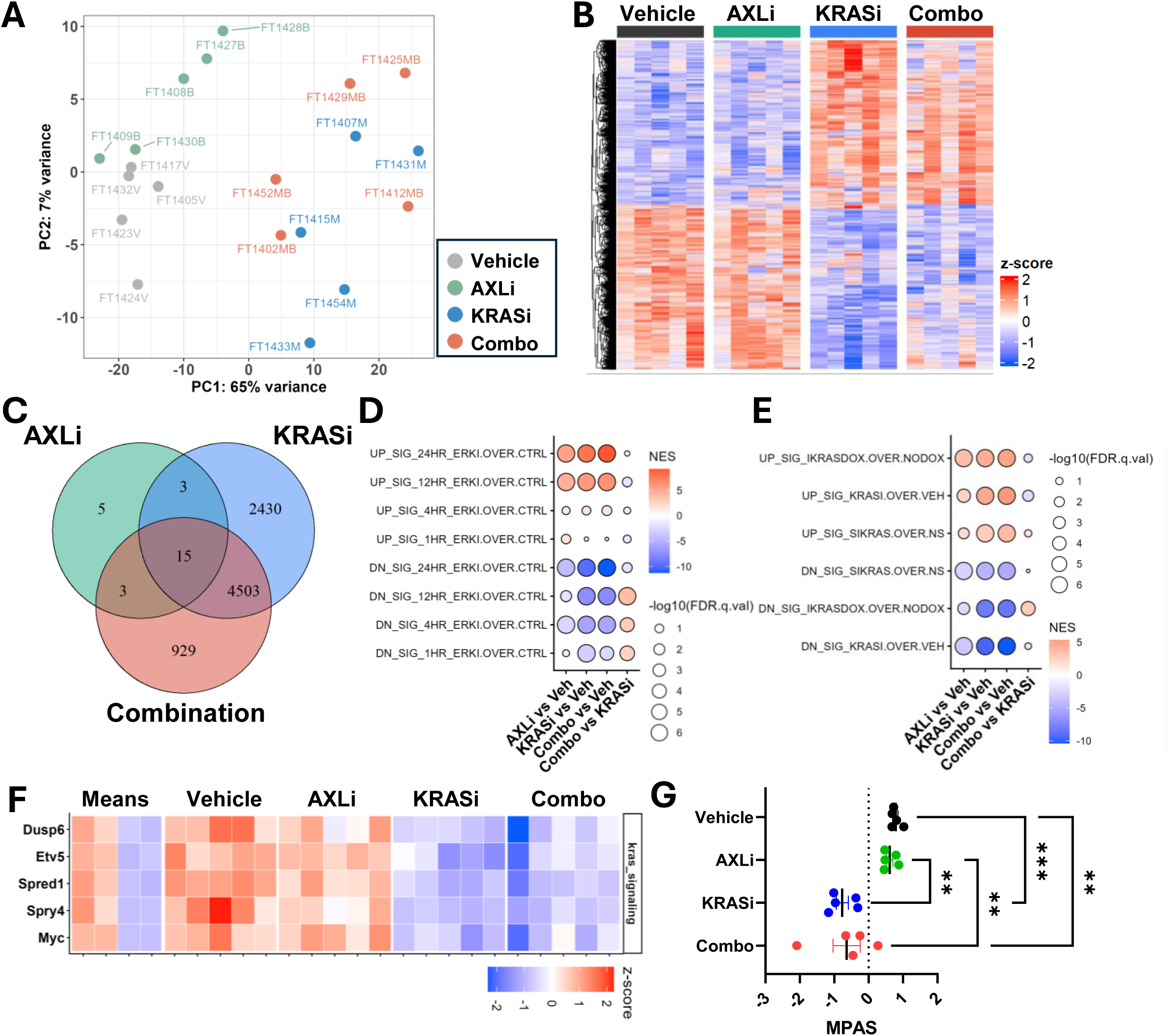
Transcriptomic analysis of KRAS and AXL inhibitor-treated tumors. Supplemental for. Figure 4. A) Subcutaneous KP1 tumor mRNA-seq sample PCA plot. B) Heatmap of all differentially expressed genes (relative to vehicle). C) Venn diagram of differentially expressed genes relative to vehicle. D) GSEA of KRAS inhibition associated gene set, E) GSEA of ERK inhibition associated gene sets. F) Tile plot of KRAS/MAPK-associated gene expression in KRASi/AXLi-treated tumors. G) MAPK Pathway Activity Score (MPAS) in tumors as function of treatment.

**Figure S7.**
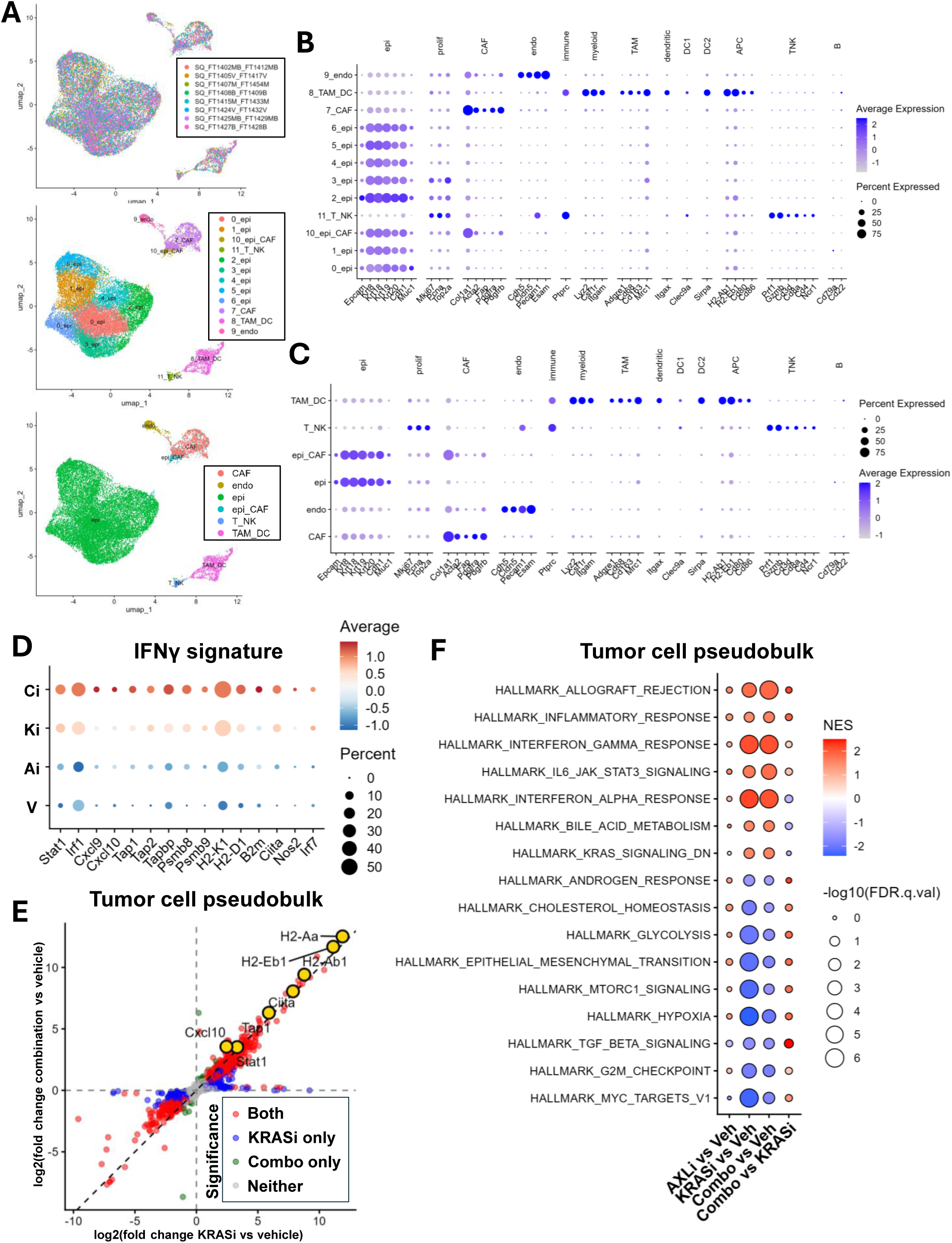
Single-cell profiling reveals tumor-intrinsic transcriptional remodeling. Supplemental for. Figure 5. A) UMAP plots of tumor samples showing sample distribution (top), Seurat clustering (middle), and broad cell-type clusters (bottom). B) Dot plot of cluster marker-expression. C) Dot plot of cell-type cluster marker-expression. D) Dot plot of IFNγ regulated genes used to calculate IFNγ module score, E) Expression fold-change vs vehicle of combination and KRAS inhibitor treated tumor gene expression in the tumor/epithelial cell cluster from pseudobulk analysis. F) GSEA of Hallmark gene sets in the tumor/epithelial cell cluster from pseudobulk analysis.

**Figure S8.**
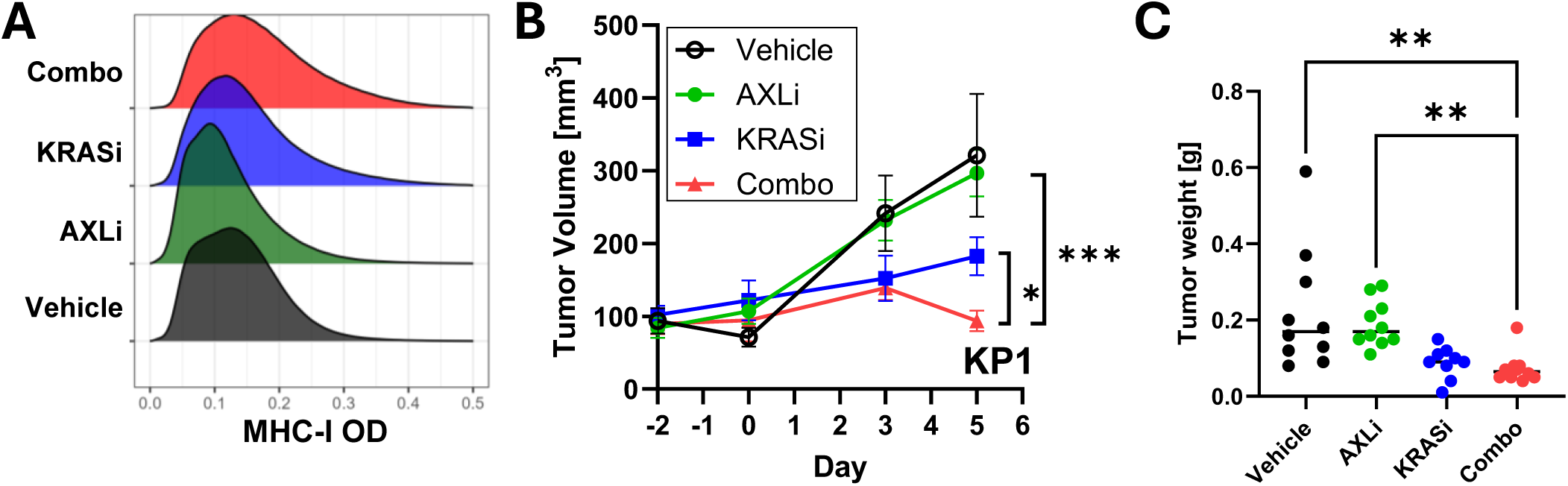
Combined KRAS and AXL inhibition enhances tumor control in vivo. Supplemental for. Figure 6. A) Distribution of MHC class I staining intensity in subcutaneous tumors as function of treatment group. B) Subcutaneous KP1 tumor volume as function of time in mice treated with KRASi (MRTX1133), AXLi, alone or in combination, or with vehicle used for immune cell profiling. C) Tumor weight as function of treatment in immune profiling cohort.

**Figure S9.**
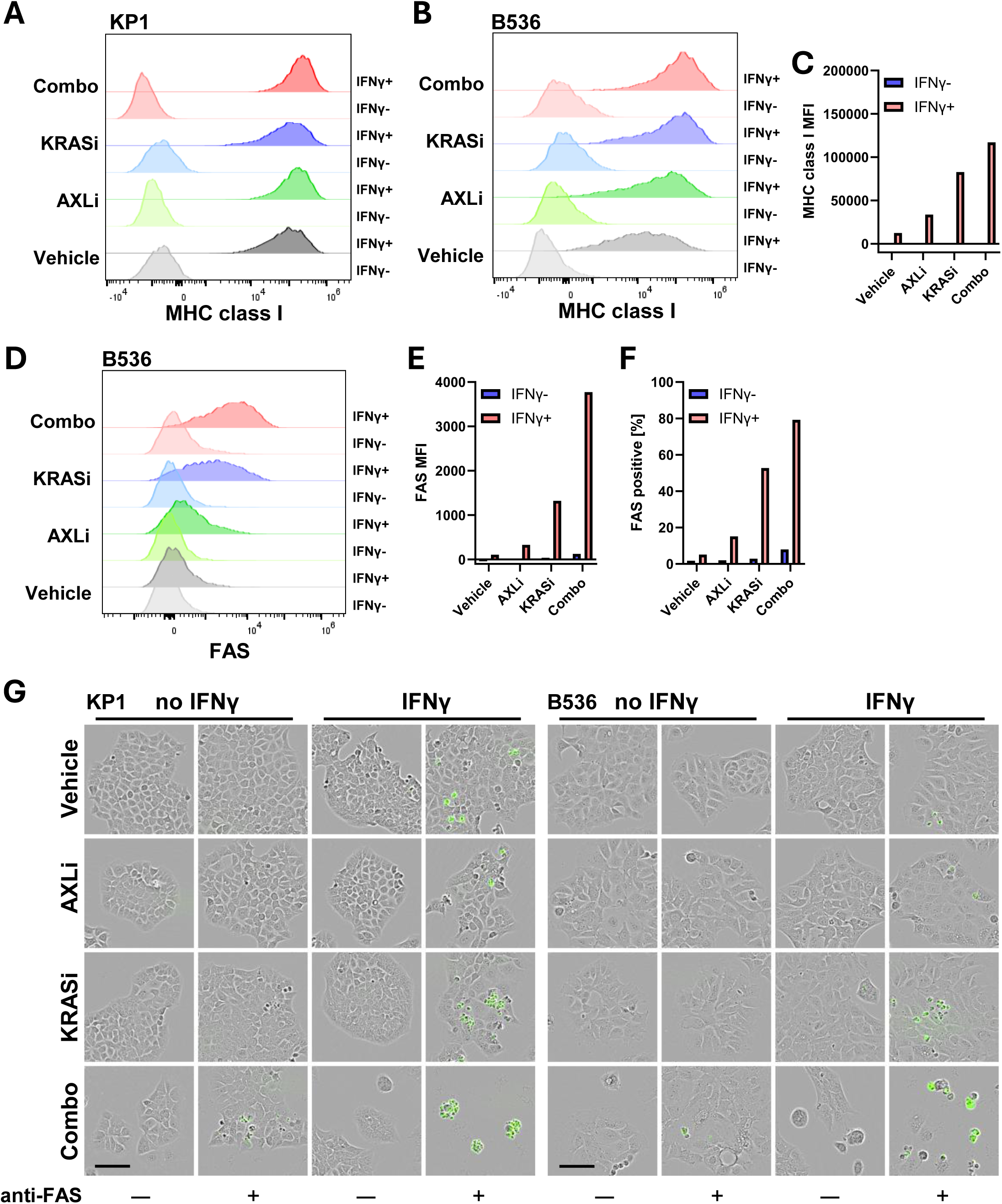
Combined KRAS and AXL inhibition sensitizes tumor cells to IFNγ and FAS stimulation. Supplemental for. Figure 7. A) Flow cytometry analysis of MHC class I surface presentation in KRASi/AXLi and IFNγ-treated KP1 cells, B) Flow cytometry analysis of MHC class I in B536 cells, C) MHC class I surface presentation (median fluorescent intensity; MFI) as function of treatment in B536 cells. D) Flow cytometry histograms of FAS surface protein presentation in B536 cells treated with KRASi/AXLi, with and without IFNγ, E) FAS surface presentation (median fluorescent intensity; MFI) in B536. F) Fraction FAS+ cells as function of treatment in B536 cells, G) Fluorescence microscopy of KP1 (left) and B536 (right) cells treated with KRASi/AXLi, IFNγ and/or agonistic FAS antibody, fluorogenic Caspase activity label in green. Scale bar 100 um.

